# Beyond One-Size-Fits-All: Tumor Biology Influences Nanoparticle Behaviour in Cancer Models

**DOI:** 10.1101/2025.03.26.645442

**Authors:** M. Bravo, G. Solís-Fernandez, S. Krzyzowska, S. Huysecom, A. Montero-Calle, I. Van Zundert, B. Louis, R. Barderas, B. Fortuni, S. Rocha

## Abstract

Nanoparticles (NPs) are a promising tool for cancer therapy, yet few have successfully reached clinical application. Current nanomedicine development pipelines are focused on optimizing physical properties of NPs, overlooking the impact of tumor biology on their behavior. Here, we show that the same NPs exhibit distinct accumulation and penetration patterns in 3D spheroids derived from four tumor models (representative of lung, colon, breast, and cervical cancer). We uncover an inverse relationship between NP uptake and penetration: tumors with slower internalization show deeper NP diffusion. Proteomic analysis revealed that tumor-specific expression of endocytic and extracellular matrix proteins underlies this variability. Our findings challenge the prevailing ‘one-size-fits-all’ approach and highlight the need to integrate tumor biology into NP design. Tailoring NPs to the unique cellular and extracellular features of each tumor type will be critical for developing more effective and clinically relevant nanotherapies.

---

Nanoparticles (NPs) have emerged as a promising alternative to conventional cancer therapies, offering unique advantages such as improved tumor targeting, localized drug delivery, and reduced systemic toxicity^1,2^. Yet, clinical translation remains limited, partly due to inconsistencies in preclinical testing that undermine the reliability of current screening approaches^2^.

Traditionally, NP development and screening rely on two-dimensional (2D) tumor models that fail to replicate key aspects of tumor physiology, masking critical biological variability between tumors^1,2^. The introduction of three-dimensional (3D) tumor models, particularly multicellular tumor spheroids, has provided a more physiologically relevant platform to assess NP accumulation, penetration, and therapeutic efficacy, while maintaining compatibility with high-throughput screening^3–5^. However, even with these improved models, a “one-size-fits-all” approach persists, with most NP development pipelines focused on tuning physical properties – such as size, shape, and surface charge – while overlooking the influence of tumour biology on NP behaviour. As a result, most studies evaluate NPs in a single tumor model and assume that findings are applicable across different cancers. However, tumor-specific features, including cell morphology, proliferation rates, vesicular trafficking and extracellular matrix (ECM) composition, can influence NP uptake and diffusion^6–8^.

This oversight has led to apparently conflicting results in the field. For instance, there is an ongoing debate concerning the effect of NP charge on spheroid penetration: Wang *et al*. reported that cationic PEGylated NPs exhibited higher penetration depth and Sujai *et al*. found that anionic PEGylated gold NPs achieved enhanced penetration^9,10^. These findings, while at first glance contradictory, were obtained in different tumor models – breast and cervical cancer, respectively – and highlight how tumor biology critically affects NP-tumor interactions, underscoring the limitations of generalizing results across tumor types. Understanding how tumor-specific characteristics shape NP behavior will be key for designing more effective, tailored NP-based cancer therapies.

In this report, we investigated how tumor biology influences NP behavior using four non-metastatic tumor models representative of cancers with high global incidence: lung (A549), colon (KM12C), breast (MCF7), and cervical (HeLa)^2,11–13^. Lung and colorectal cancers rank amongst the most commonly diagnosed worldwide, while breast and cervical cancer are highly prevalent among women^11^. As a model NP system, we used gold nanospheres encapsulated within a mesoporous silica shell, which was subsequently coated with polyethyleneimine (PEI, Au@mSi-PEI NPs, Fig. S1). The silica shell improved biocompatibility and dispersion in aqueous media, while the outer cationic PEI layer enhanced membrane interactions and NP uptake^14,15^. The inherent photoluminescence of the gold core under multiphoton excitation enabled direct NP monitoring via fluorescence microscopy^2,16^.

## Tumor-Specific Differences in Nanoparticle Distribution in 3D Models

To assess how tumor biology affects NP behavior in solid tumors, we analyzed the accumulation and distribution of Au@mSi-PEI NPs in the four different cancer models. Spheroids were incubated for 3, 6, 24, and 48 hours to capture the temporal dynamics of NP transport within the tumor models.

Clear differences in NP accumulation were observed across tumor types (Fig. 1a, 1b, S2, S3). At early time points (3 and 6 hours), overall accumulation was low in all spheroids, although KM12C and MCF7 spheroids exhibited slightly higher NP levels compared to A549 and HeLa. By 24 hours, NP accumulation more than doubled in all models, with KM12C showing the highest accumulation, and A549 the lowest. After 48 hours, KM12C spheroids maintained the highest NP levels, while HeLa exhibited the lowest accumulation. Notably, HeLa spheroids, which had relatively high NP levels at 24 hours, exhibited a marked drop by 48 h. This coincided with a decrease in spheroid size (Fig. S5), suggesting that the proliferative outer layer, which contained most of the NPs, detached and was lost during sample handling. As a result, NP accumulation and penetration data in HeLa spheroids became less reliable at this time point.

**Figure 1.**
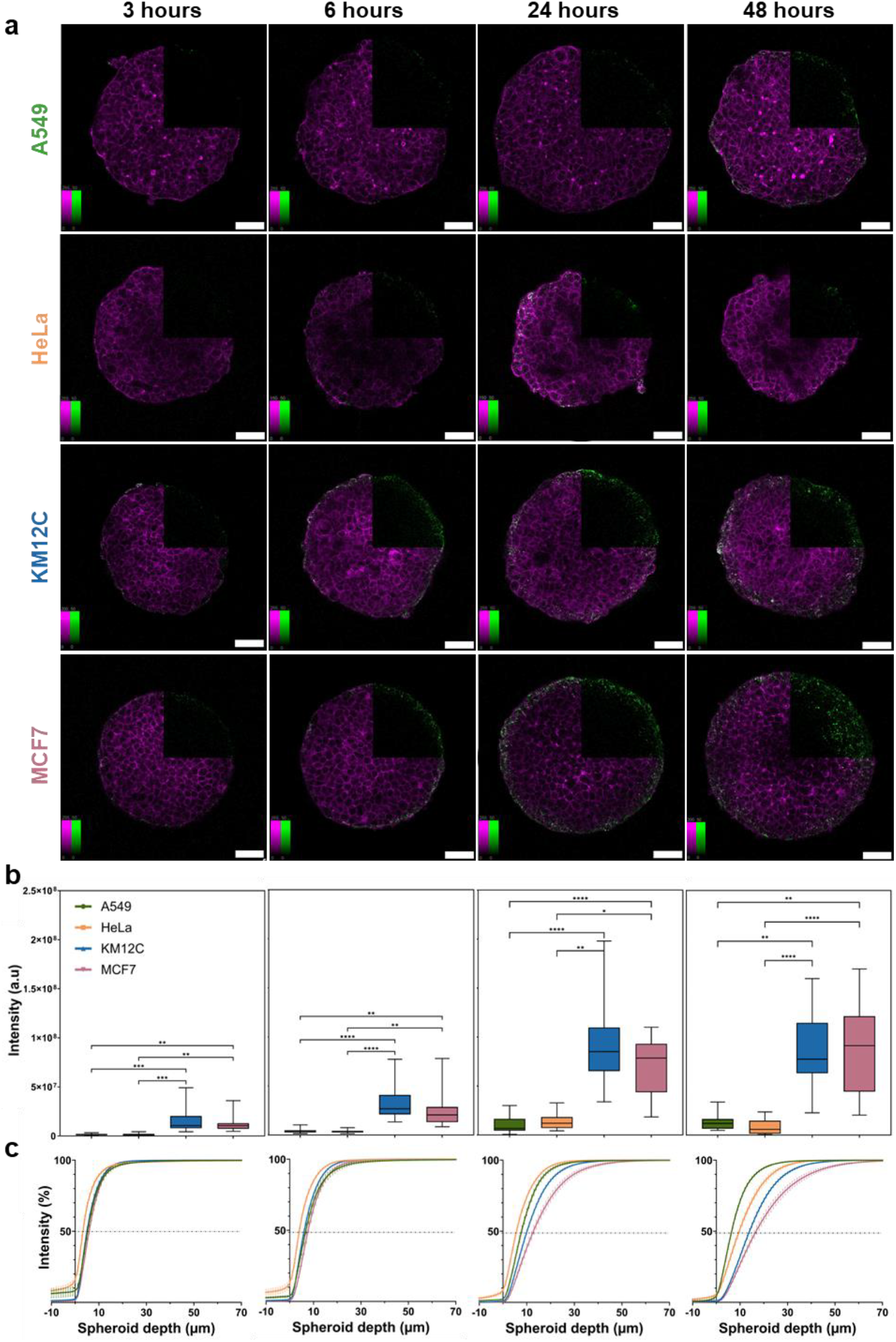
(a) Confocal fluorescence microscopy images of NP distribution in 3D tumor spheroids from A549 (lung), HeLa (cervical), KM12C (colorectal), and MCF7 (breast) cancer cell lines, at the mid-plane of the spheroids (with a diameter of approximately 250 µm). Au@mSi-PEI distribution was assessed at 3, 6, 24, and 48 hours, with columns showing the different time points and rows corresponding to the cell lines. Nanoparticles (green) were detected via photoluminescence following multiphoton laser excitation, and actin filaments (magenta) were stained with Phalloidin CruzFluor^TM^ 647. Scale bar = 50 µm. Color bars indicate contrast values, fixed for the NP channel (0–50). Additional images are provided in Supplementary Figure S2. (b) Total NP accumulation per spheroid, based on total NP signal intensity within the spheroid volume. (c) NP penetration profiles displayed as cumulative NP distribution (intensity %) relative to the distance from the spheroid outer rim at each time point. A549, HeLa, KM12C, and MCF7 are represented in green, orange, blue, and pink, respectively. Error bars indicate ± SD, with ns meaning not significant; * (p < 0.05), ** (p < 0.01), *** (p < 0.001), and **** (p < 0.0001) (n ≥ 20).

NP penetration profiles followed a similar time-dependent trend. At 3 and 6 hours, NP distribution was restricted to the outer cell layers, with 95% of the NPs (P95) found within the first 23 µm of the spheroids (Fig. 1a, 1c, Fig. S2, S4). By 24 hours, MCF7 and KM12C spheroids exhibited deeper penetration, with P95 values of 36 µm and 27 µm, respectively, while NPs in A549 and HeLa spheroids remained confined to a smaller region (P95 between 22 and 19 µm). After 48 hours, MCF7 spheroids showed the greatest penetration depth, with 5% of the NPs beyond 47 µm, followed by KM12C spheroids (P95 of 34 µm). In contrast, NP penetration in A549 and HeLa spheroids remain low (P95 of 21 and 28 µm, respectively).

These findings reveal tumor-dependent differences in NP accumulation and penetration within 3D spheroids, suggesting that tumor-specific factors govern NP distribution in solid tumors. To investigate underlying mechanisms driving this variability, we evaluated NP-cell interactions at the single-cell level.

### NP Uptake Dynamics Across Tumor Models

While 3D spheroids better mimic tumor physiology, their size and complexity hinder the quantification of NP uptake within individual cells. To address this limitation, we used 2D monolayers as a complementary model to precisely monitor subcellular NP localization. 2D cellular systems also enable pulse-chase experiments, where NPs are incubated for a defined period and then removed to track intracellular NP dynamics over time – an approach less feasible in spheroids due to the difficulty of removing extracellular NPs without disturbing the 3D structure. Here, cells were incubated with Au@mSi-PEI NPs for 3 h, after which NPs were removed. Imaging was performed immediately (defined as 3 hours) and at 6, 24, and 48 hours after NPs addition (Fig. 2a), to monitor NP uptake and retention over time.

**Figure 2.**
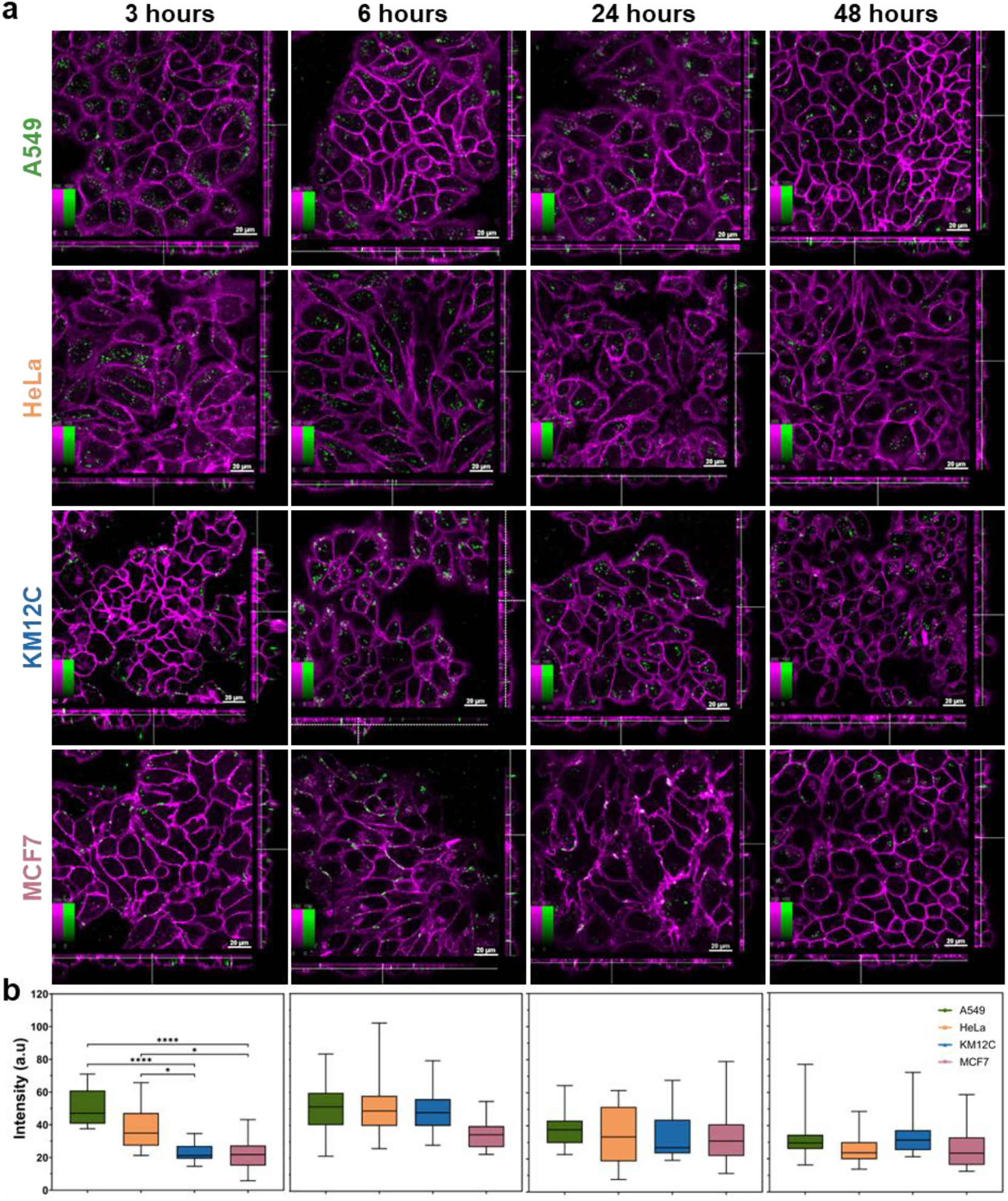
(a) Confocal fluorescence microscopy images of NP internalization in 2D cell monolayers from A549 (lung), HeLa (cervical), KM12C (colorectal), and MCF7 (breast) cancer cell lines. AuNP@mSi@PEI uptake was assessed at 3, 6, 24, and 48 hours, with columns showing the different time points and rows representing cell lines. Nanoparticles (green) were detected via photoluminescence following multi-photon laser excitation, and cell membrane (magenta) was stained with CellMask^TM^ DeepRed. Scale bar = 20 µm. Color bars show contrast values, fixed for the NP channel (0–100). Additional images are provided in Supplementary Figure S5. The central square represents a single xy plane, while the bottom and left panels are the xz and yz cross-sections (b) Average NP intensity inside the cells, normalized to the cell area. A549, HeLa, KM12C, and MCF7 are represented in green, orange, blue, and pink, respectively. Error bars indicate ± SD, with ns meaning not significant; * (p < 0.05), ** (p < 0.01), *** (p < 0.001), and **** (p < 0.00010 (n ≥ 20).

NP uptake kinetics varied across tumor models, displaying distinct trends over time (Fig. 2b, S6, S7). While A549 and HeLa internalized substantial amounts of NPs within the first 3 hours, KM12C and MCF7 displayed minimal uptake. By 6 hours, NP uptake in HeLa and KM12C cells increased, reaching levels comparable to A549. MCF7 consistently exhibited the lowest internalization, with NPs largely remaining at the cell membrane rather than within the cytoplasm. At later time points (24 and 48 hours), NP accumulation became more uniform across cell lines, showing no significant differences.

These findings show that NP uptake dynamics vary significantly across tumor cell lines and inversely correlate with NP penetration in 3D spheroids. For example, rapid NP uptake by A549 cells correlated with NP retention in the spheroid’s outer layers. In contrast, MCF7 displayed the slowest NP uptake in monolayers, which was associated with the deepest NP penetration in spheroids (Fig. 1c, 2b). This suggests that rapid NP uptake by peripheral cells may act as a trap, limiting further diffusion into deeper layers. Conversely, slower uptake in KM12C and MCF7 cells likely enables NPs to travel further before being internalized, potentially via paracellular transport.

### Inverse Relationship Between Nanoparticle Uptake and Penetration in Tumor Spheroids

To test whether NP uptake and tumor penetration are indeed inversely related, we evaluated the behavior of NPs lacking the PEI coating. Bare silica-shell gold NPs, which possess a negative surface charge (Au@mSi, Fig. S1), are expected to show reduced membrane interaction and lower internalization^14,15^.

In agreement with previous reports, removal of the PEI coating reduced NP internalization and resulted in a more uniform uptake across all tumor models in 2D monolayers (Fig. 3a, b, S9)^14,15^. This decrease in cellular uptake translated into enhanced NP penetration in 3D spheroids (Fig. 3c, d). At 48 hours, penetration depths increased across most tumor models: A549 spheroids, which previously showed highly restricted diffusion, exhibited the largest improvement (P95 increasing from 21 µm to 42 µm). KM12C and MCF7 spheroids also showed moderate increases in penetration (P95 from 34 µm to 45 µm, and from 47 µm to 50 µm, respectively). In contrast, HeLa spheroids showed no increase in the P95 value, likely due to the loss of the outer proliferative rim, which compromised spheroid integrity and affected penetration analysis.

**Figure 3.**
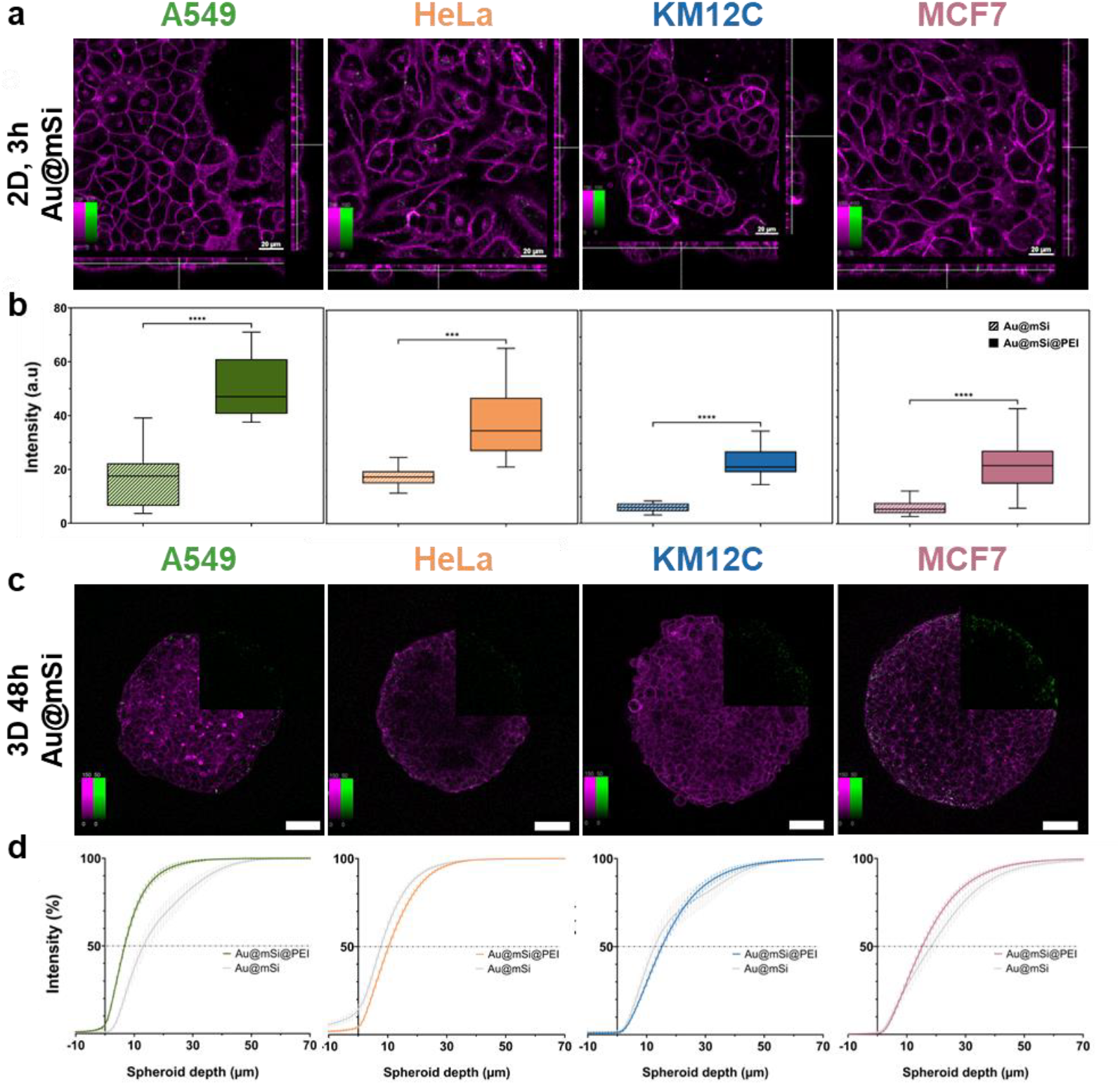
(a) Confocal fluorescence microscopy images of Au@mSi internalization in 2D cell monolayers of A549 (lung), HeLa (cervical), KM12C (colorectal), and MCF7 (breast) cancer cell lines. AuNP@mSi uptake was assessed at 3 hours. Nanoparticles (green) were detected via photoluminescence following multi-photon laser excitation, and cell membrane (magenta) was stained with CellMask^TM^ DeepRed. Scale bar = 20 µm. (b) Average NP intensity inside the cells, normalized to the cell area. For each cell line, NP uptake for Au@mSi and Au@mSi-PEI is compared. (c) Confocal fluorescence microscopy images of NP distribution in 3D tumor spheroids from A549, HeLa, KM12C, and MCF7 cancer cell lines, at the mid-plane of the spheroids (with a diameter of approximately 250 µm). Au@mSi distribution was assessed at 48 hours. Nanoparticles (green) were detected via photoluminescence following multiphoton laser excitation, and actin filaments (magenta) were stained with Phalloidin CruzFluor^TM^ 647. Scale bar = 50 µm. Additional images are provided in Supplementary Figure S8. (d) NP penetration profiles displayed as cumulative NP distribution (Intensity %) relative to the distance from the spheroid outer rim, comparing NP distribution of Au@mSi-PEI and Au@mSi NPs for each cell line at 48 hours. A549, HeLa, KM12C, and MCF7 are represented in green, orange, blue, and pink, respectively. Error bars indicate ± SD, with ns meaning not significant; * (p < 0.05), ** (p < 0.01), *** (p < 0.001), and **** (p < 0.00010 (n ≥ 20).

These findings confirm that limiting NP uptake at the spheroid periphery promotes deeper NP penetration, supporting the inverse correlation between cellular uptake and NP diffusion.

### Tumor Proteomics and Nanoparticle Uptake and Penetration

Given the key role of endocytosis in NP internalization, we hypothesized that the inverse relationship between NP uptake and penetration is driven by tumor-specific differences in the expression of endocytic proteins.

Quantitative proteomic analysis was performed across the four tumor models, resulting on the identification of 7236 proteins. To focus on endocytosis, we further analyzed proteins classified under the gene ontology (GO) term “Endocytic Transport” (GO: 0030139). Principal component analysis (PCA) showed high reproducibility across the 4 replicates and revealed that KM12C and MCF7 spheroids exhibited similar expression profiles of endocytic proteins, distinct from both A549 and HeLa (Fig. 4a). This mirrors the NP uptake profiles, where breast and colorectal cancer cells displayed slower internalization compared to HeLa and A549.

**Figure 4.**
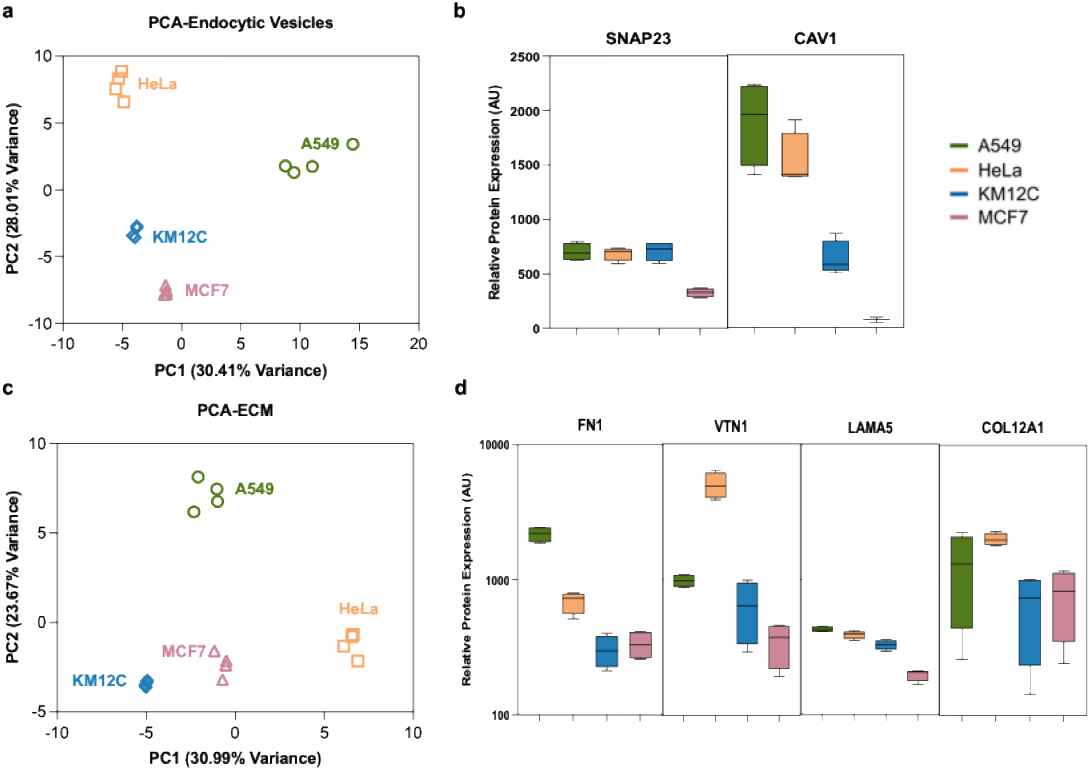
(a) Principal component analysis (PCA) of Endocytosis -related proteins across four tumor spheroid models. PCA was performed on quantitative proteomic data from four biological replicates per condition, and the four points inside each cluster representing the four biological replicates used for proteomics analysis. (b) Protein expression levels of two of the most relevant endocytic proteins (SNAP23 and CAV1). (c) Principal component analysis (PCA) of ECM -related proteins across four tumor spheroid models. PCA was performed on quantitative proteomic data from four biological replicates. (d) Protein expression levels of four of the most relevant ECM proteins (COL1A1, FN1, VTN, and LGALS1). A549, HeLa, KM12C, and MCF7 are represented in green, orange, blue, and pink, respectively. Error bars indicate ± SD, with ns meaning not significant; * (p < 0.05), ** (p < 0.01), *** (p < 0.001), and **** (p < 0.00010 (n ≥ 4).

To pinpoint candidate regulators of NP uptake, we analyzed proteins with significant changes in expression levels compared to MCF7 cells, that showed the highest NP penetration and lowest uptake (fold-change ≥ 2 or ≤ 0.5; Table S1, Fig. 4b). Among the identified proteins, the synaptosome-associated protein-23 (SNAP23), essential for membrane fusion and early endosomal function, was upregulated in A549 (2.1-fold), HeLa (2.1-fold), and KM12C cells (2.2-fold), suggesting improved cellular trafficking in these tumors^17^. Furthermore, the CAV1 gene, a scaffolding protein associated with caveolae-mediated endocytosis, showed overexpression in A549 (25.7-fold), HeLa (20.8-fold), and KM12C (8.6-fold) spheroids, correlating with the uptake trend observed in 2D monolayers. This suggests that caveolae-dependent mechanisms are a key driver of Au@mSi-PEI NP internalization in these tumor models.

In addition to endocytic pathways, we investigated whether tumor extracellular matrix (ECM) composition also contributes to the distinct NP behavior observed across tumor models. Previous studies have shown that ECM properties can influence NP behavior, particularly in solid tumor models^18,19^. Using GO classification under the term “Extracellular Matrix” (GO: 0031012), we identified tumor-specific differences in the expression of ECM-related proteins (Table S2, Fig. S4c, d). In comparison to MCF7 spheroids, A549 and HeLa spheroids – which exhibited limited NP penetration – showed higher expression of key structural ECM components associated with increased matrix density and stiffness, including fibronectin (FN1), vitronectin (VTN), laminin alpha subunit 5 (LAMA5), and collagen alpha-1(XII) chain (COL12A1). These findings suggest that a denser ECM may act as an additional physical barrier to NP diffusion, complementing the role of cellular uptake in modulating NP transport.

## Conclusions

Our results show that tumor-intrinsic characteristics affect NP behavior, challenging the current ‘one-size-fits-all’ paradigm in NP design. Distinct tumor types displayed unique NP accumulation and penetration profiles in 3D spheroids, driven by differences in endocytosis-related protein expression and extracellular matrix (ECM) composition. Using 2D monolayers, we uncovered an inverse relationship between NP internalization and diffusion depth, where tumors with slower NP uptake allowed for deeper NP penetration, likely via paracellular routes. This hypothesis was supported by the enhanced penetration of NPs without PEI. Proteomic analysis confirmed that higher expression of endocytosis-related proteins aligned with increased NP internalization, while ECM-stiffening proteins were upregulated in tumor models showing reduced NP diffusion.

These findings highlight the need to move beyond solely optimizing NP physicochemical properties and instead tailor designs to tumor-specific biological features. Moreover, therapeutic goals should guide optimization strategies: drug delivery applications may benefit from high NP uptake, whereas therapies like photothermal treatments may require deeper NP diffusion into the tumor mass. By integrating tumor biology into NP development, therapeutic efficacy can be improved, ultimately accelerating the clinical translation of nanomedicines.

## Materials and Methods

### Materials

Dulbecco’s modified eagle medium (DMEM), gentamicin, phosphate buffered saline (PBS, no calcium, no magnesium), paraformaldehyde (v/v 16%, methanol-free), Trypsin-ethylenediaminetetraacetic acid (Tryspin-EDTA, 0.5% solution, no phenol red), Hank’s balanced salt solution (HBSS, no phenol red), Triton X-100 (0.1%), UltraPure™ agarose (2%), sodium chloride (NaCl), GlutaMax^TM^ , Trypan blue solution (0.4%, TC grade), and CellMask^TM^ DeepRed were purchased from **ThermoFisher Scientific**. Tetrachloroauric(III) acid trihydrate (HAuCl4 · 3H2O, 99%), sodium hydroxide (NaOH, 98%), hexadecyltrimethylammonium bromide (CTAB, ≥99%), tetraethyl orthosilicate (TEOS, 98%), hydrochloric acid (HCl, 1 N), methanol (MeOH, 99,8%), polyethyleneimine solution (PEI, 50% w/v in H_2_O), 3D Petri Dish® micro-mold, fetal bovine serum (FBS), and CoverWell^TM^ perfusion chambers (9 mm × 1.7 mm thickness) were purchased from **Sigma Aldrich**. Phalloidin CruzFluor^TM^ 647 was purchased from **Santa Cruz Biotechnology**. A549 (lung cancer), MCF-7 (breast cancer) cell lines were a gift from Prof. Uji-i. All chemicals were used without further purifications.

## Methods

### Nanoparticle Synthesis and Functionalization

Mesoporous silica-coated gold nanoparticles (Au@mSi) were synthesized using a one-pot method as described by Chen *et al*^20^. Initially, 50 mg of CTAB was dissolved in a solution containing 0.6 mL of 0.5 M NaOH and 24 mL of Mili-Q water. The mixture was stirred at 800 rpm and 80°C for 15 minutes. Subsequently, 1 mL of a 2% (wt) aqueous formaldehyde solution was added, followed by 0.8 mL of a 0.05 M HAuCl_4_ aqueous solution. After 10 min, a solution of TEOS and ethanol (1 mL, containing 0.25 g TEOS and 0.50 g ethanol) was added, and the mixture was stirred for an additional hour. The resulting product was collected by centrifugation, dispersed in a 1.1 M HCl solution in water/ethanol (v/v = 1.25:10), and sonicated. The dispersion was then stirred vigorously at 60°C for 4 h to remove CTAB from the pores. The product was washed twice with Milli-Q water to neutralize the pH, followed by redispersion in 3.2 mL of Milli-Q water. The mixture was stirred magnetically at 500 rpm for 3 h, and the supernatant was removed by centrifugation and replaced with Milli-Q water. For PEI functionalization, a 0.75% (w/v) PEI solution (adjusted to pH 7) was added dropwise to the Au@mSi NPs in a 1:1 (v/v) ratio. The mixture was stirred for 3 h, washed by centrifugation, and redispersed in Milli-Q water. The concentration of particles in the colloidal solution was estimated to be 1.3 mg/mL.

### AuNP@mSi-PEI Characterization

The physicochemical properties of the NPs were characterized using UV-Vis spectroscopy, scanning and transmission electron microscopy (SEM, TEM), Zeta Potential, and Dynamic Light Scattering (DLS). UV-Vis spectra were acquired using a Cary 60 Spectrophotometer (Agilent). High-resolution SEM images were captured with a field-emission SEM (FEI Quanta FEG250) microscope operated at 20.0 kV. For TEM measurements, NPs were diluted in a 1:100 ratio, and 6 μL of solution was cast onto TEM grids (300 mesh copper grid) and dried at room temperature. TEM images were collected on an FEI TecnaiF20 microscope (200 kV) (Ian Holmes Imaging Center, Bio21). Particle size, size distribution, polydispersity index, and Zeta Potential measurements were performed using a Zetasizer Nano ZS (Malvern Panalytical). All measurements were carried out in Milli-Q water at room temperature (25°C).

### Cell Culture

A549, HeLa, KM12C, and MCF7 cells were cultured in 25 cm^2^ culture flasks at 37° C under a 5% CO_2_ atmosphere. Cells were maintained in culture medium (DMEM supplemented with 10% FBS, 1% L-glutamax, and 0.1% gentamicin), and passaged using trypsin-EDTA at 80-90% confluency.

### NP Screening in Spheroids (3D)

At 90% confluency, cells were harvested for spheroid preparation. Spheroids were cultured in agarose microtissues (16 x 16 array of 256 wells), cast from ultra-pure agarose (20 mg/mL in Milli-Q water with 9 mg/mL NaCl) using a 3D micro-mold (3D Petri Dish®, Sigma Aldrich). Once solidified, the agarose microtissue was removed from the mold, placed in a sample dish, and equilibrated with DMEM for 15 min. After removing the DMEM, 190 μL of cell suspension was added to the mold. Cell concentrations were optimized to achieve spheroids of comparable sizes: A549 and MCF7 were seeded at 1.3×10^6^ cell/mL, HeLa cells at 1.6×10^6^ cell/mL, and KM12C at 2.5×10^5^ cell/mL. Cells were allowed to sink into the agarose wells for 30 min, followed by the addition of DMEM around the microtissue. Samples were incubated for 4 days to obtain fully formed spheroids before incubation with NPs.

Once spheroids were fully formed, the DMEM inside and around the microtissues was removed. A 200 μL suspension of Au@mSi-PEI NPs or Au@mSi (26 µg/mL) in DMEM was added to the microtissues and incubated for 30 min. DMEM was then added around the microtissues to prevent dilution of the NP suspension. To assess NP accumulation and penetration, spheroids were incubated with Au@mSi-PEI for 3, 6, 24, and 48h. Au@mSi assays were only performed for the 48h time point. Following incubation, the medium was removed and 1 mL of PBS was vigorously pipetted onto the microtissue to dislodge the spheroids, which were transferred to 1.5 mL microcentrifuge tubes.

Harvested spheroids were stained with Phalloidin CruzFluor^TM^ 647^21^. Spheroids were washed three times with PBS, and collected using 10-second centrifugation (2000 G). They were then fixed with 500 μL 4% PFA for 20 min, followed by three PBS washes. Permeabilization was performed with 1 mL 0.1% Triton X-100 for 30 min. Spheroids were incubated overnight at room temperature with 500 μL of Phalloidin CruzFluor^TM^ 647 (1:800 dilution in 3%BSA), for cytoskeleton (actin) staining. The next day, the spheroids were washed 3 times with PBS and resuspended in 200 μL of PBS before imaging. For spheroid imaging, 60 μL of spheroid suspension was transferred to a CoverWell^TM^ perfusion chamber glued on a #1 glass coverslip.

### NP Screening in Cell Monolayers (2D)

Cells (2×10^5^ cells/dish), were seeded in 29-mm glass-bottom dishes (Cellvis, Mountain View, CA, USA) and incubated for 24 hours, to reach 60-80% confluency. NP uptake was assessed at 3, 6, 24, and 48h. Cells were incubated with Au@mSi-PEI NPs (26 µg/mL) for 3 hours, then carefully washed three times with PBS. For the 3-hour time point, cells were immediately stained and imaged. For subsequent time points, fresh DMEM was added, and samples were incubated at 37°C for the remaining time. Before imaging, the cell membrane was stained with CellMask™ Deep Red (1 μM in HBSS, 5 min), washed three times with PBS, and imaged in HBSS.

### Fluorescence Microscopy

Confocal fluorescence imaging of 2D and 3D samples was performed on a Leica TCS SP8 dive inverted microscope. This system is equipped with a multi-photon (MP) Insight X3 laser (range 680-1300 nm), two HyD detectors (non-de-scanned), a 25× water immersion objective (NA 0.95, FLUOTAR VISIR) for 3D models, and a 63× oil immersion objective (NA 1.4) for 2D models. Sequential scanning (between stacks) was performed, starting with the Phalloidin CruzFluor 647^TM^ channel (1200 nm, 3.8 mW at the objective, 610-661 nm detection), followed by the Au@mSi/Au@mSi-PEI NP channel (800 nm, 3.8 mW at the objective, 503-553 nm detection). For 2D imaging, live imaging was performed in HBSS at 37°C and 5% CO_2_. In this case, the first sequence targeted the CellMask^TM^ Deep Red channel using a 638 nm diode laser (detection 641-691 nm). Image stacks of 1024 × 1024 pixels were acquired at a scanning speed of 400 Hz, 0.57 or 0.30 µm z-step, for 3D and 2D assays, respectively.

### NP Behavior Analysis

In-house algorithms were used to determine NP accumulation/penetration in 3D spheroids and uptake in 2D cell monolayers. NP distribution profiles in 3D were obtained from the analysis of at least 20 spheroids. For 3D spheroids, the algorithm first preprocessed cytoskeleton-staining images of the spheroid to identify the spheroid edge, by applying smoothing, dilation and edge detection. The spheroid edge was then used to obtain the spheroid center and a spherical coordinate system. This coordinate system enabled the calculation of edge-to-center distance at each angle of elevation (θ) and azimuth (ψ). For the NP channel, the same spherical coordinates were used to calculate the NP penetration depth by measuring the distance of the NPs from the spheroid edge at the same angles θ and ψ. Integration of pixel intensities across penetration depths provided individual penetration profiles, which were then averaged to obtain the mean distribution profile (Fig. S10).

NP behavior in 2D was analyzed from at least 20 images of different fields of view. The algorithm was used to quantify the total intensity of the NPs in the cytoplasm. First, cytoplasm and cellular membrane were segmented based on the membrane staining intensity by using the Cellpose algorithm in MatLab^22^. NPs were quantified by integrating the fluorescence intensity signal of the NP signal within each cell volume (Fig. S11).

### LC-MS/MS and proteomic analysis

Cell pellets of spheroids from all the cell lines in the absence or presence of NPs (24h incubation following the protocol described in methods section NP Screening in Spheroids (3D)) were collected via centrifugation. The cell pellets wheren then lysated with RIPA buffer (Sigma Aldrich) containing 1X protease and phosphatase inhibitors (MedChemExpress). Total protein concentration was obtained by the tryptophan quantification method as described in (REF) and confirmed by coomassie blue staining after reducing 10% PAGE-SDS.

Once protein extracts from spheroids in the presence or absence of NPs were obtained, 10 µg of each extract were reduced, alkylated, and trypsin digested on SP3 magnetic beads as previously described^23^.

For LC-MS/MS, peptides were analyzed in an Orbitrap Astral mass spectrometer coupled to a Vanquish Neo UHPLC System (Thermo Fisher Scientific). Peptide samples were loaded into the precolumn PepMap Trap Cartridge 5 µm, 300 µm x 5 mm (Thermo Fisher Scientific) and eluted in an Easy-Spray PepMap RSLC C18 3 µm, 75 µm x 15 cm (Thermo Fisher Scientific) heated at 50°C. The mobile phase flow rate was 300 nL/min and 0.1% formic acid (FA) in H2Omq and 0.1% FA in 80% acetonitrile (ACN) were used as elution buffers A and B, respectively. The 15 min elution gradient was: 4%-10% buffer B for 2 min, 10%-40% buffer B for 11 min, 40%-99% buffer B for 0.5 min, and 99% buffer B for 1.5 min. Prior to injection, samples were re-suspended in 20 µL of buffer A, and 2 µL of each sample were injected, and analyzed in data independent acquisition (DIA) mode. For ionization, 1900 V of liquid junction voltage and 280°C capillary temperature was used. The full scan method employed a m/z 380-980 mass selection, an Orbitrap resolution of 240000 (at m/z 200), an automatic gain control (AGC) value of 500%, and maximum injection time (IT) 5 ms. The MS/MS was performed with the Astral mass analyzer, using an AGC of 500%, an IT of 3 ms, and a normalized collision energy (NCE) of 25 for fragmentation of precursors. The scan range was set from 380 to 980 m/z, with an isolation window of 2 m/z, and window placement optimization was enabled. Thus, a total of 299 windows were analyzed in each cycle.

Raw data were analyzed with Spectronaut (version 19.1.240724.62635) using standardized workflows. DIA raw data and Uniprot UP000005640_9606.fasta Homo sapiens (march, 2024) database (20,418 protein entries) were used for the construction of the spectral library by directDIA. Trypsin/P and Lys/P were selected as the digestion enzymes and a maximum of 2 missed cleavages were allowed. Carbamidomethylation of cysteines was set as a fixed modification, and methionine oxidation and N-terminal acetylation were set as variable modifications. For DIA analysis, standard workflow was used. The maximum FDR for peptide spectral match (PSM), peptide, and protein identifications was set at 0.01, automatic cross-run normalization was enabled, and all identified peptides were used for protein quantification. Protein inference was performed using the IDPicker algorithm. Non imputation was performed during peptide and protein identification and quantification with Spectronaut.

### Gene Ontology (GO) analysis

The list of proteins identified through proteomic analysis were filtered to obtain which components of the GO terms “Extracellular Matrix” (GO: 0031012, 1826 annotations), “Endocytic Vesicle” (GO: 0030139, 958 annotations) and “Exocytic Vesicle” (GO: 0070382, 549 annotations) in homo sapiens were present in the cells under study. Each list of filtered proteins was then used to perform Principal Component Analysis (PCA) based on the expression levels of the proteins. PCA plots were obtained using Python and sklearn.decomposition and sklearn.preprocessing packages. The list of filtered proteins after removing repeated annotations can be found in the supplementary tables S2-4.

## Acknowledgements

The author(s) declare that financial support was received for the research, authorship, and/or publication of this article. MB acknowledges the Global PhD partnership program between KU Leuven and Melbourne University (GPUM/21/025). We acknowledge additional financial support from Research Foundation of Flanders (FWO) research grants (G0D4519N, G081916N, VS08523N, G0C1821N, and G022724N), postdoctoral fellowships (for BF: 12X1419N and 12X1423N, for IV: 12A6N25N, for BL: 12AGZ24N and for GSF: 12AML24N), and from the KU Leuven (IDN/20/021, C14/15/053, C14/19/079, and C14/22/085). This work also received financial support from PI20CIII/00019 and PI23CIII/00027 grants from the AES-ISCIII program cofounded by FEDER funds to R.B.

## Supplementary Information

### Nanoparticle Synthesis and Characterization

Au@mSi NPs were prepared as previously described by Chen *et al*^20^. Gold nanospheres (AuNPs) with a mesoporous silica layer (Au@mSi) in a core-shell structure were synthesized using a one-pot method. A PEI layer (Mw = 1.3 kDa) was then added to the Au@mSi surface, through electrostatic interactions^15^. Transmission electron microscopy (TEM) and scanning electron microscopy (SEM) revealed spherical NPs with homogeneous size distribution and minimal aggregation. AuNP cores exhibited an average diameter of 43 nm, while the mSi shell was approximately 15 nm thick, resulting in Au@mSi of (77 ± 8) nm (Fig. S1a-c).

The localized surface plasmon resonance (LSPR) was characterized using UV-VIS spectroscopy (Fig S1c), showing a redshift of the plasmonic band after mSi coating, due to an increase in the local refractive index^24^. Zeta potential measurements showed that AuNP cores (Au@CTAB) initially displayed a positive charge of (36 ± 1) mV due to the presence of CTAB (Cetylmethylammonium bromide) on the NP surface. To enhance aqueous stability, the mSi layer was first added, followed by the removal of CTAB, which reduced toxicity^25^. During this process, CTAB acted as a pore-generating agent, forming micelles that serve as templates for silica condensation^26^. The resulting Au@mSi displayed a negative charge (- 30 ± 1 mV), as a result of the deprotonated hydroxyl groups on the silica surface^15^. Lastly, upon PEI functionalization, the zeta potential values increased significantly (+ 43 ± 1 mV), due to the amine groups in PEI. Due to its positive charge, PEI is expected to enhance NP interaction with the negatively charged cell membrane, promoting their uptake through endocytic pathways^14^.

**Figure S1.**
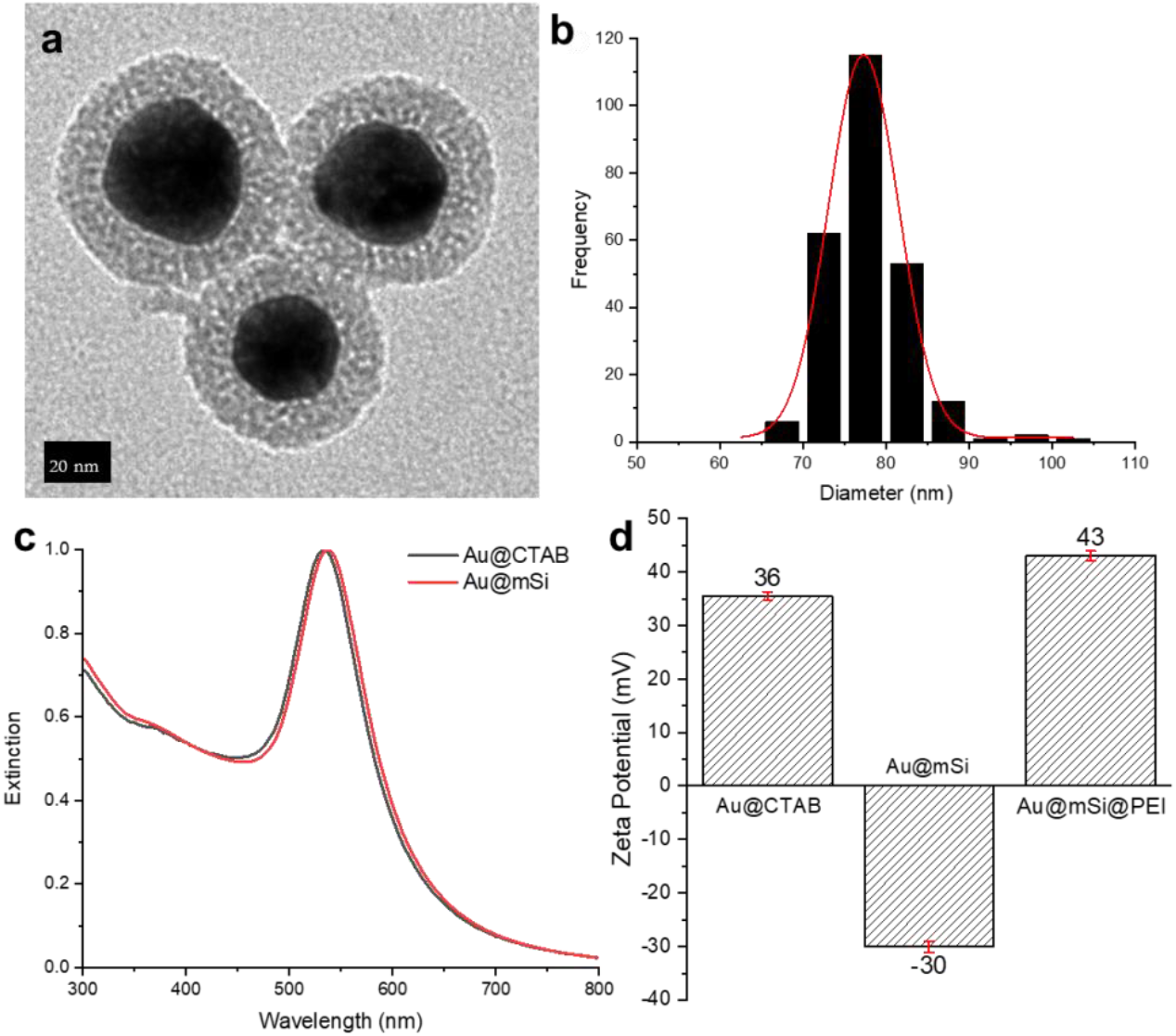
(a) Transmission electron Microscopy (TEM) image of Au@mSi NPs, showing homogeneous mesoporous silica coating of individual AuNPs. Scale bar = 20 nm; (b) Representative Transmission Electron Microscopy (TEM) image of Au@mSi NPs. Scale bar = 20 nm; (c) Size distribution plot of Au@mSi NPs; (d) Extinction coefficient of AuNP before and after mSi coating (in water); (e) Zeta potential measurements of AuNPs with different functionalizations, given as mean ± SD (n = 3).

### Nanoparticle Diffusion and Accumulation in 3D Spheroids

**Figure S2.**
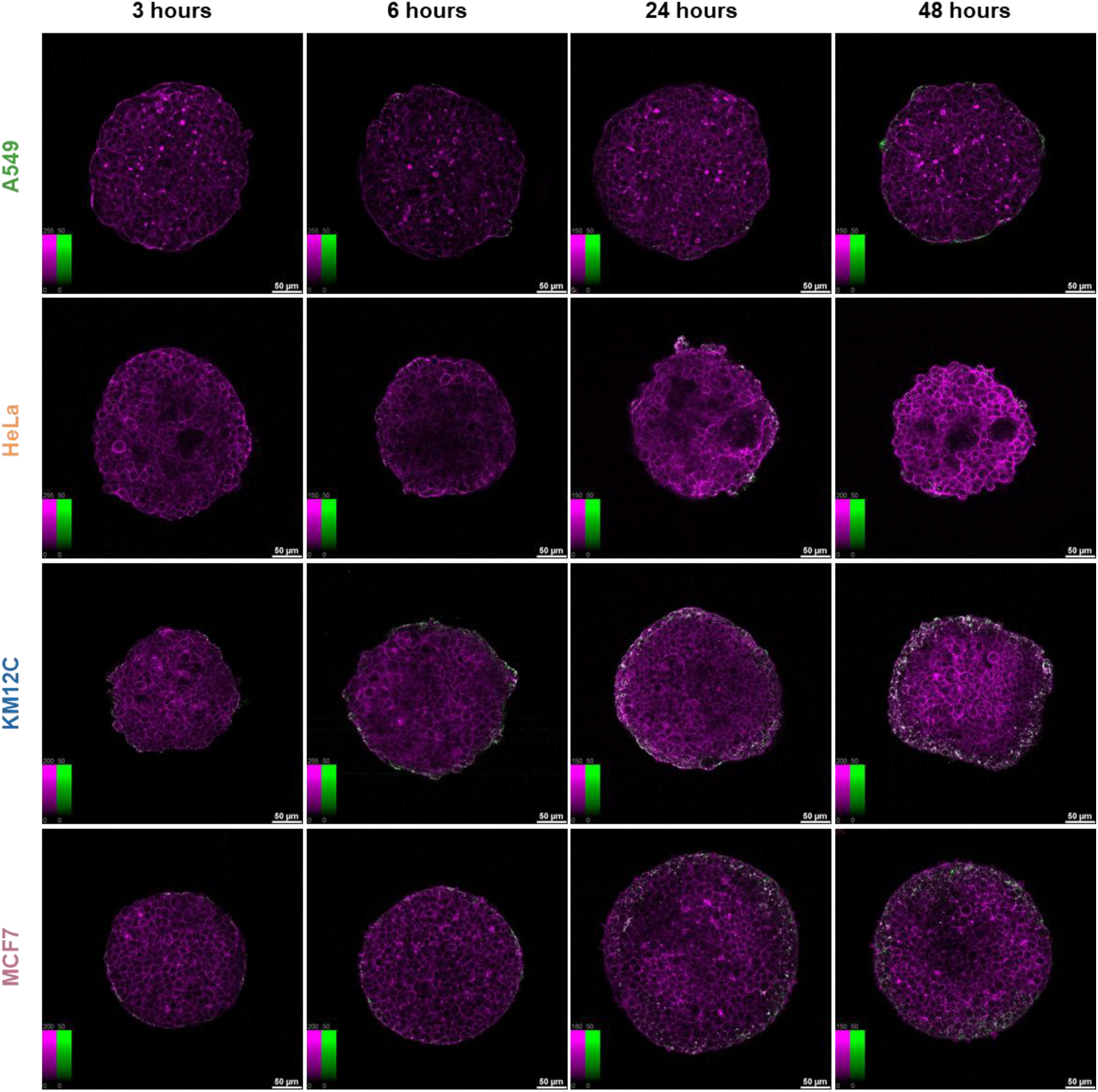
Confocal fluorescence microscopy images of NP distribution in 3D tumor spheroids from A549 (lung), HeLa (cervical), KM12C (colorectal), and MCF7 (breast) cancer cell lines, at the mid-plane of the spheroids. AuNP@mSi@PEI uptake was assessed at 3, 6, 24, and 48 hours, with columns showing the different time points and rows representing cell lines. Nanoparticles (green) were detected via photoluminescence following multi-photon laser excitation, and actin filaments (magenta) were stained with Phalloidin CruzFluor647. Scale bar = 50 µm. Color bars show contrast values, fixed for the NP channel (0–50).

**Figure S3.**
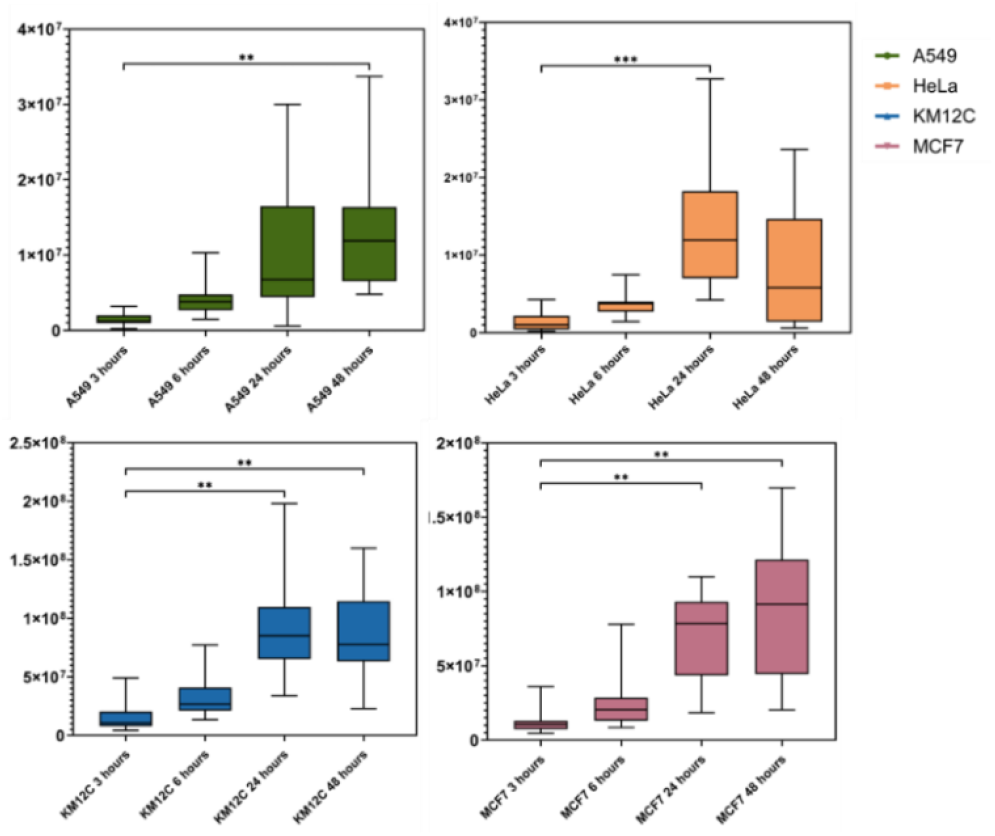
Average total NP accumulation per spheroid, based on total NP signal intensity within the spheroid volume at the different time points, showing increasing amount of NPs inside the spheroids over time for all the samples, except HeLa spheroids after 48 hours of NP incubation, where a slight intensity decrease was observed. A549, HeLa, KM12C, and MCF7 are represented in green, orange, blue, and pink, respectively. Error bars indicate ± SD, with ns meaning not significant; * (p < 0.05), ** (p < 0.01), *** (p < 0.001), and **** (p < 0.00010) (n ≥ 20).

**Figure S4.**
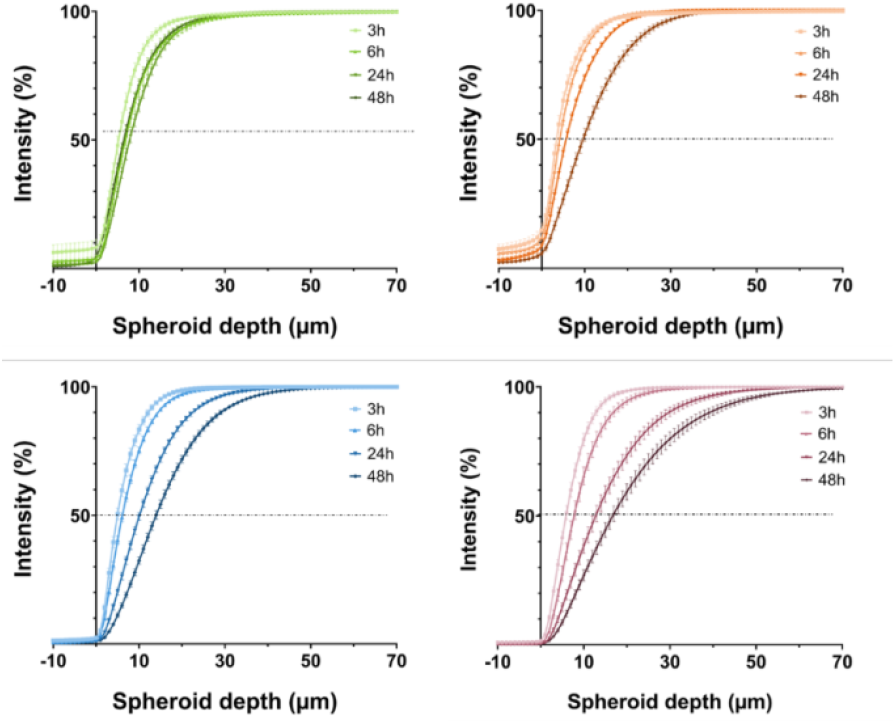
NP penetration profiles displayed as cumulative NP distribution (Intensity %) relative to the distance from the spheroid outer rim at each time point. A549, HeLa, KM12C, and MCF7 are represented in green, orange, blue, and pink, respectively. (n ≥ 20).

**Figure S5.**
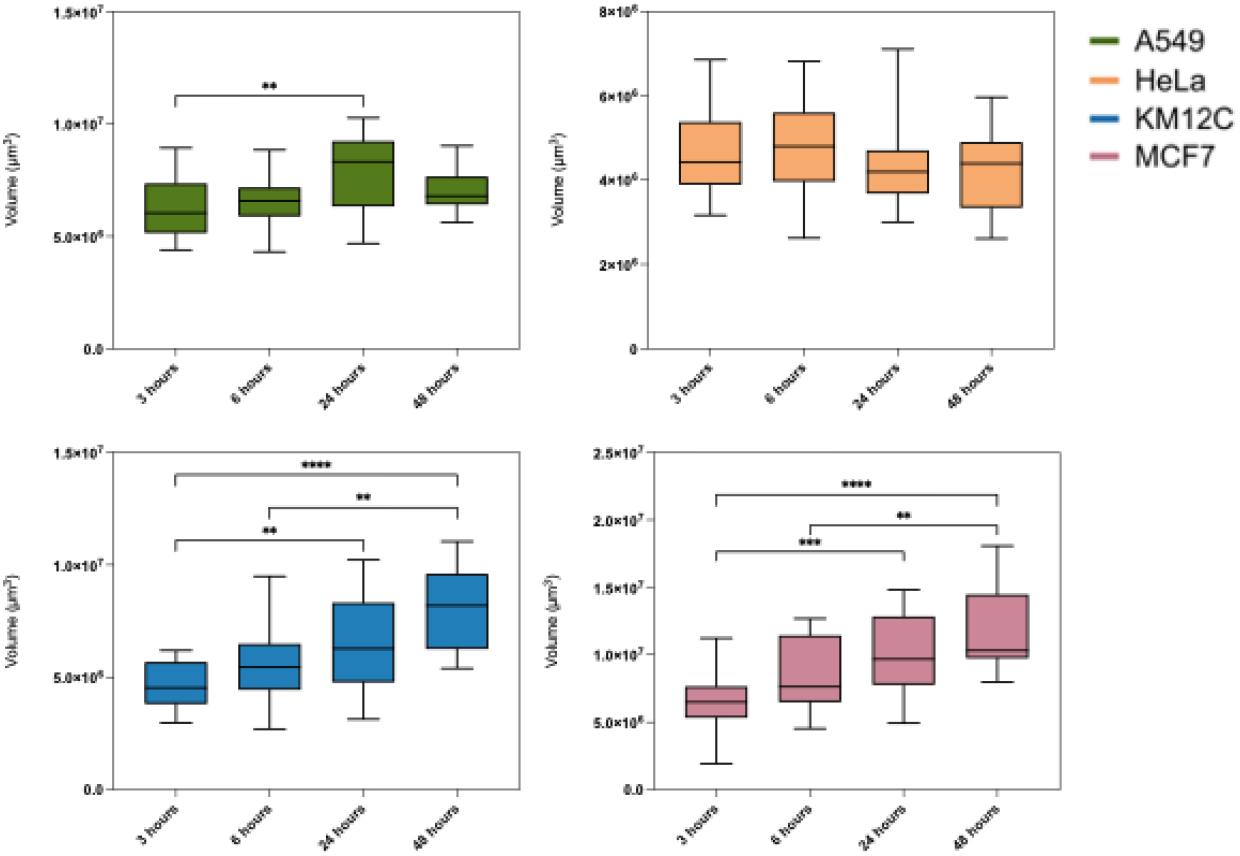
Spheroid size over time. A549, Hea, KM12C, and MCF7 are represented in green, orange, blue, and pink, respectively. (n ≥ 20). Error bars indicate ± SD, with ns meaning not significant; * (p < 0.05), ** (p < 0.01), *** (p < 0.001), and **** (p < 0.00010) (n ≥ 20).

### Nanoparticle Uptake in 2D Cell Monolayers

**Figure S6.**
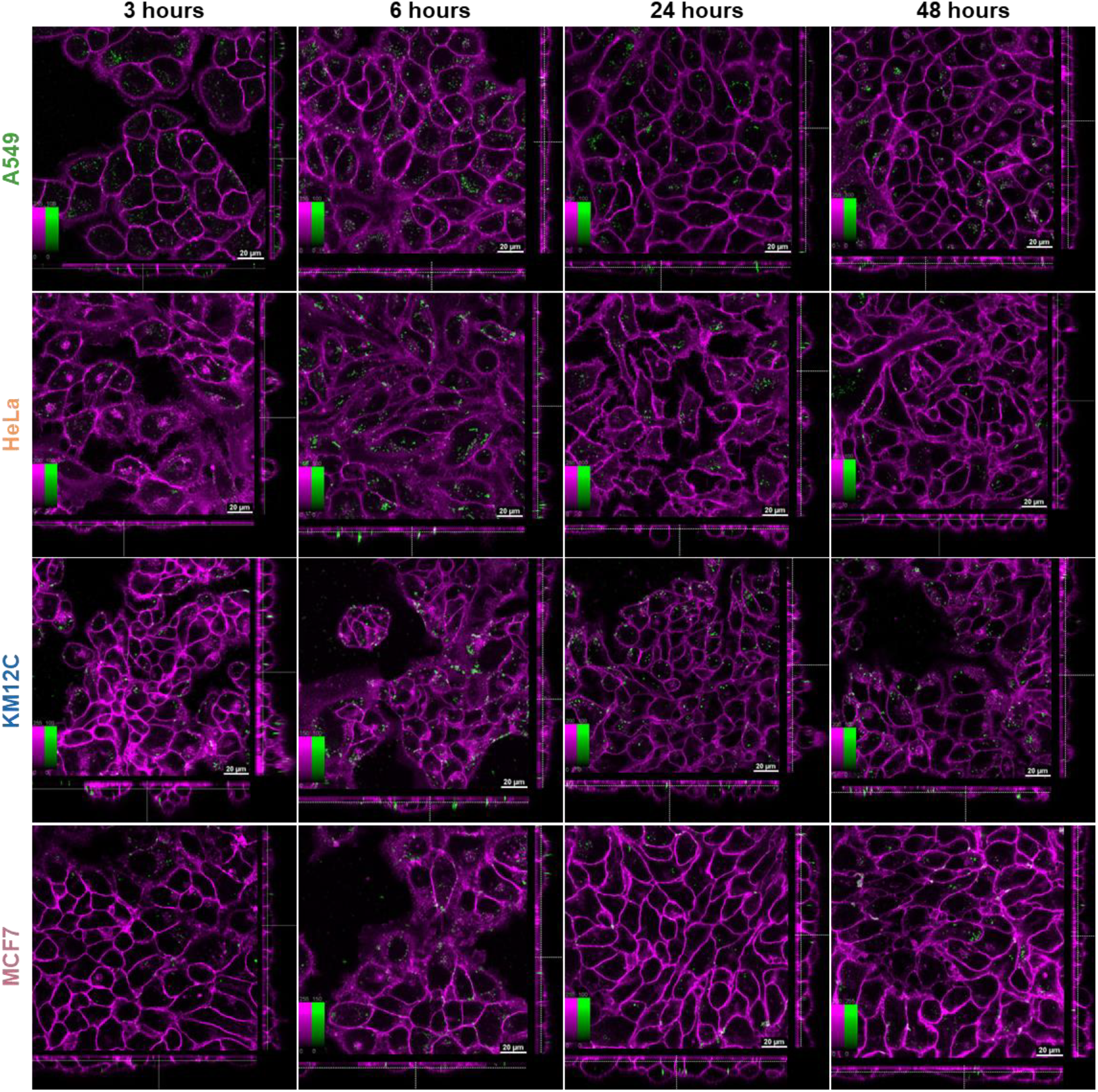
Confocal fluorescence microscopy images of NP internalization in 2D cell monolayers from A549 (lung), HeLa (cervical), KM12C (colorectal), and MCF7 (breast) cancer cell lines. AuNP@mSi@PEI uptake was assessed at 3, 6, 24, and 48 hours, with columns showing the different time points and rows representing cell lines. Nanoparticles (green) were detected via photoluminescence following multi-photon laser excitation, and cell membrane (magenta) was stained with CellMask DeepRed. Scale bar = 20 µm. Color bars show contrast values, fixed for the NP channel (0–100). The central square represents a single xy plane, while the bottom and left panels are the xz and yz cross-sections.

**Figure S7.**
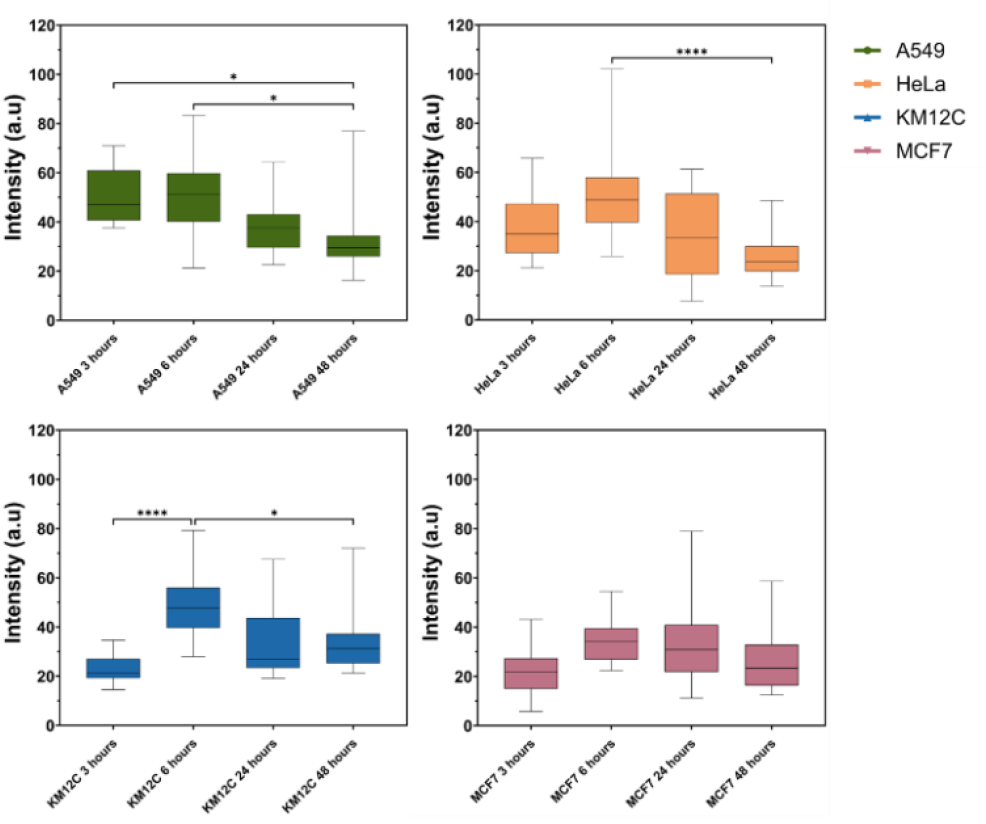
Average NP intensity inside the cells per cell line over time, normalized to the cell area. A549, HeLa, KM12C, and MCF7 are represented in green, orange, blue, and pink, respectively. Error bars indicate ± SD, with ns meaning not significant; * (p < 0.05), ** (p < 0.01), *** (p < 0.001), and **** (p < 0.00010 (n ≥ 20).

### Au@mSi NP Behavior in 2D and 3D Tumor Models

**Figure S8.**
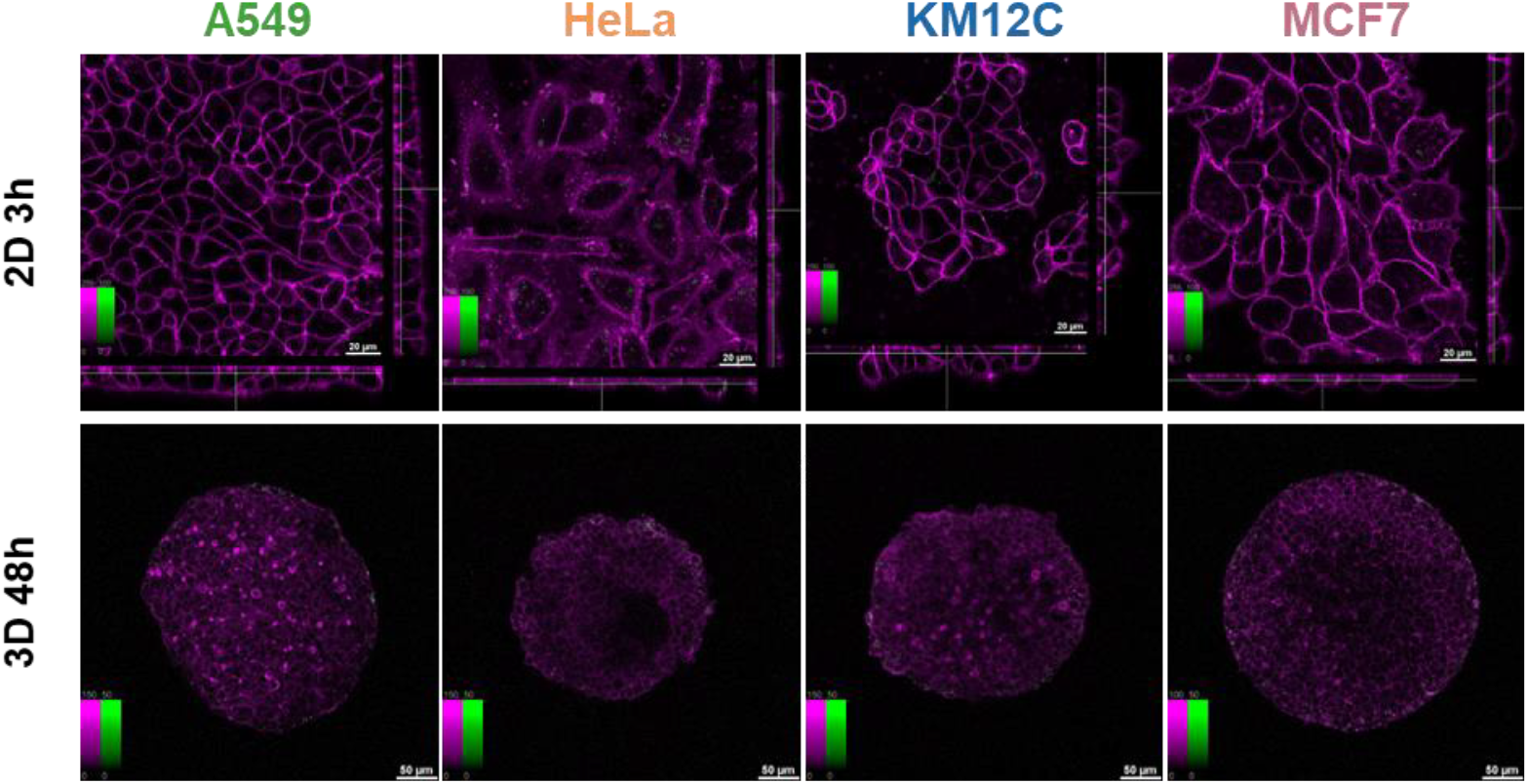
Confocal fluorescence microscopy images of Au@mSi internalization in 2D cell monolayers and 3D spheroids from A549 (lung), MCF7 (breast), HeLa (cervical), and KM12C (colorectal) cancer cell lines. AuNP@mSi uptake was assessed at 3 hours in 2D and 48 hours in 3D models, with columns showing the different cell lines. Nanoparticles (green) were detected via photoluminescence following multi-photon laser excitation, and cell membrane (2D, magenta) or cytoskeleton (3D, magenta) were stained with CellMask DeepRed and Phalloidin CruzFluor647, respectively. Scale bar: 20 µm in cell monolayers and 50 µm in tumor spheroids. Color bars show contrast values, fixed for the NP channel (0–100 in 2D, and 0-50 in 3d). The central square in 2D monolayers represents a single xy plane, while the bottom and left panels are the xz and yz cross-sections.

**Figure S9.**
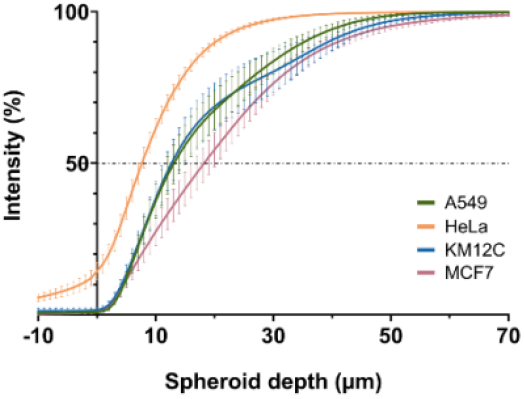
NP penetration profiles displayed as cumulative NP distribution (intensity %) relative to the distance from the spheroid outer rim, comparing NP distribution of Au@mSi NPs for each cell line at 48 hours. A549, HeLa, KM12C, and MCF7 are represented in green, orange, blue, and pink, respectively. (n ≥ 20).

### Analysis of Endocytic Protein Expression

**Table S1.**
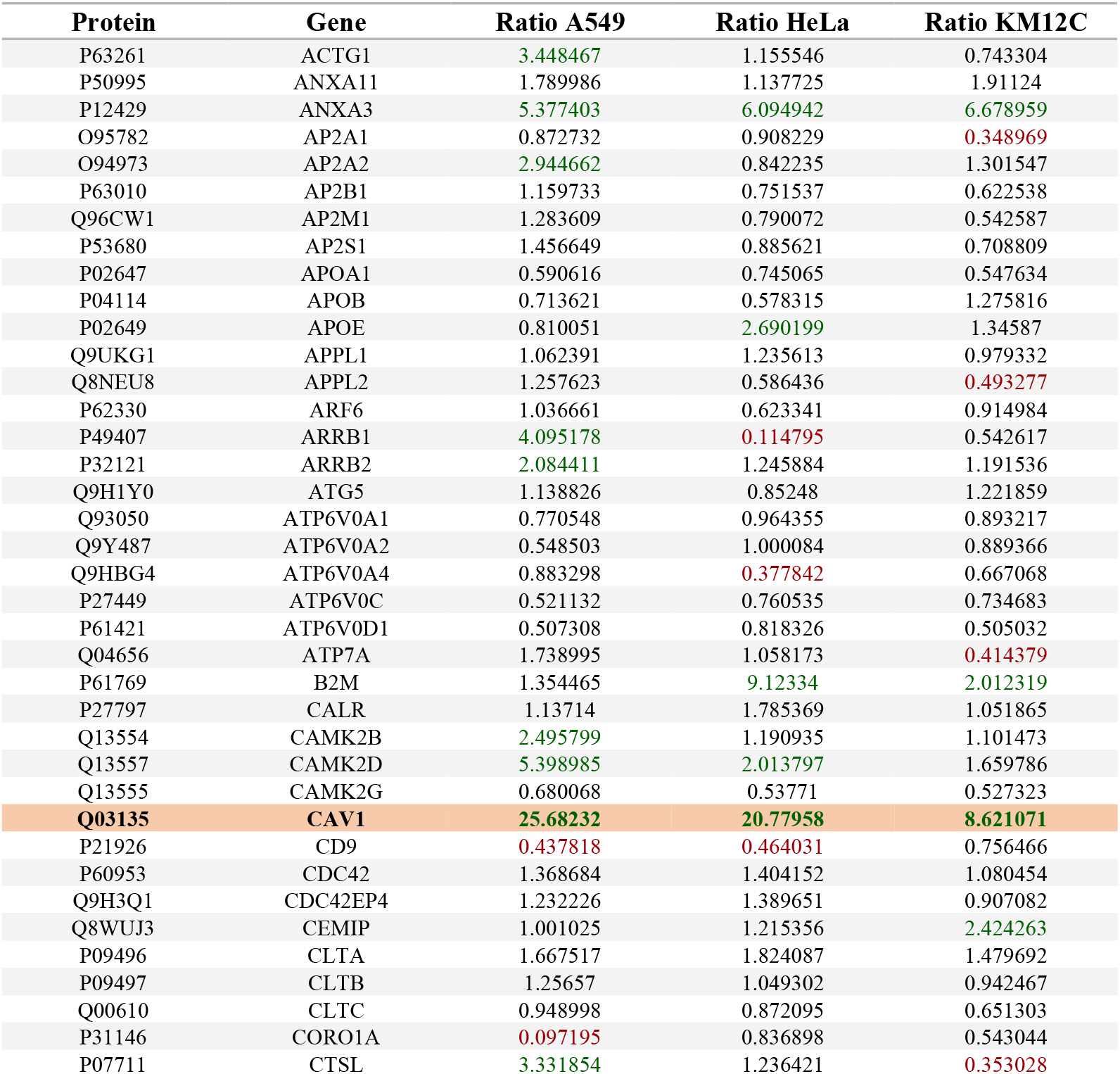
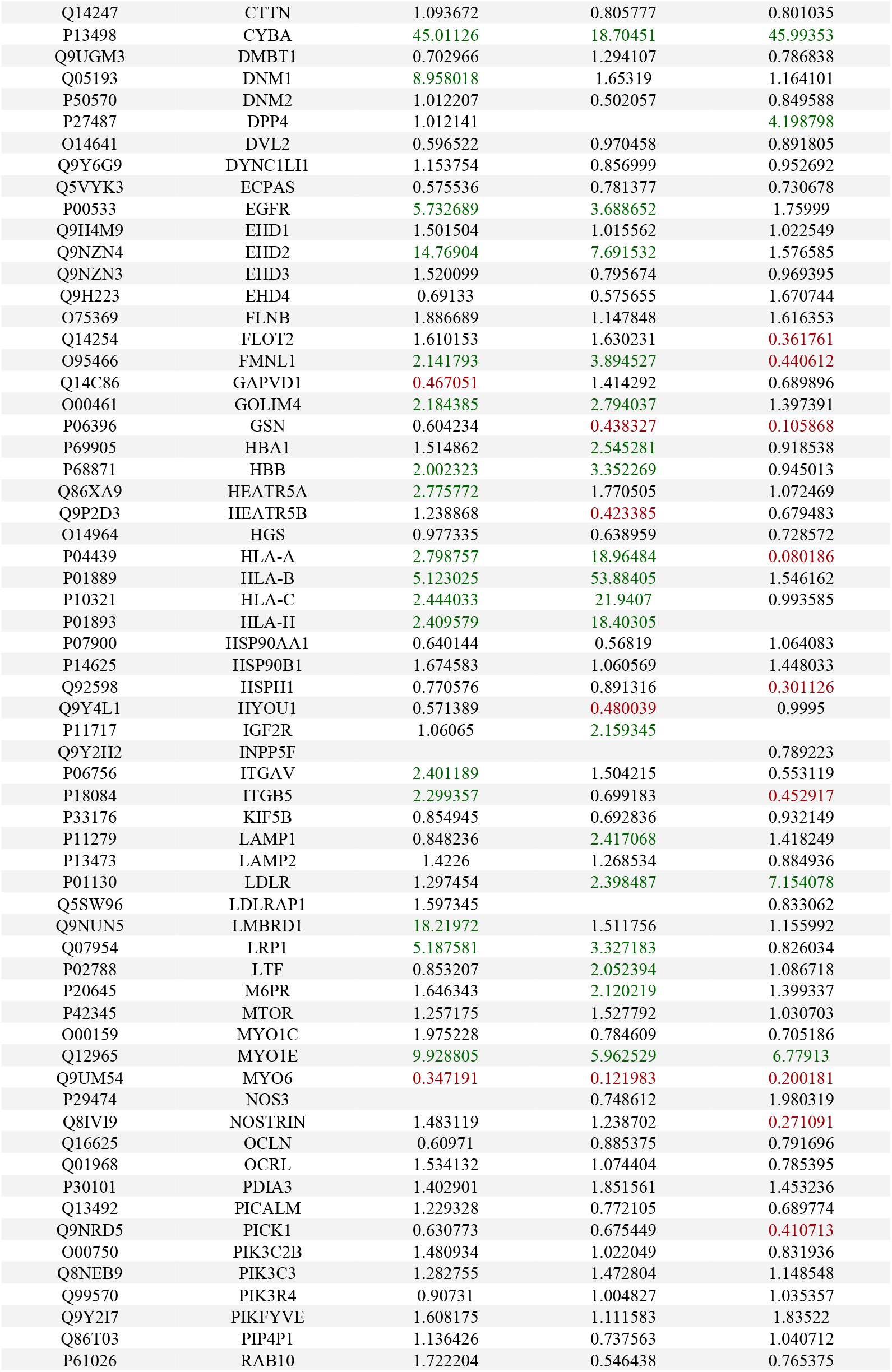
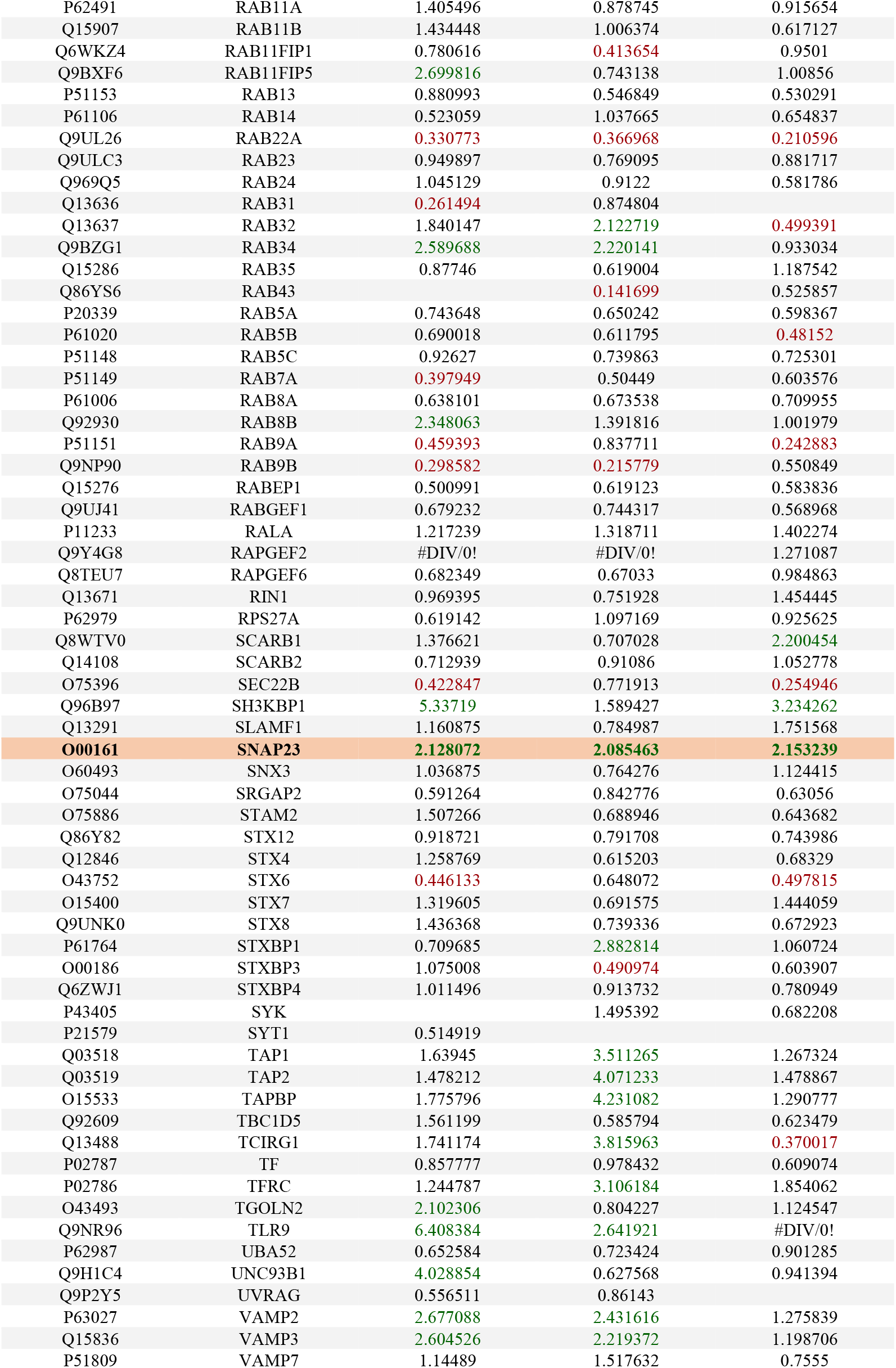
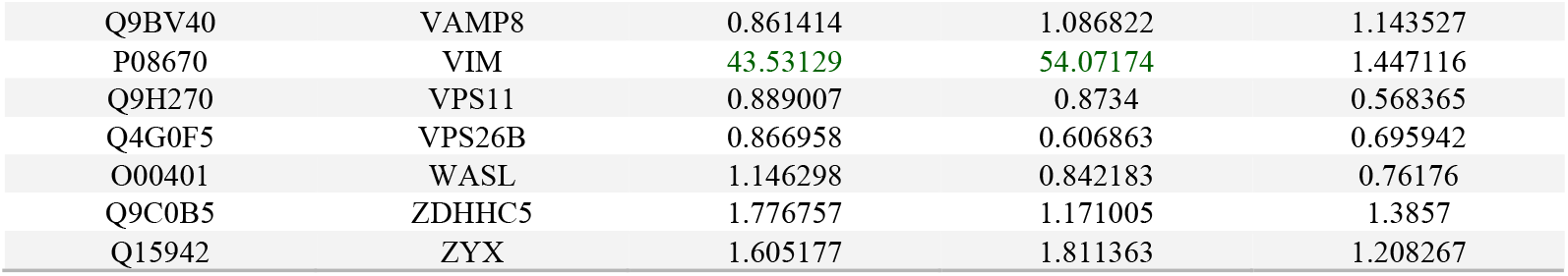
Endocytosis-related proteins, filtered by carrying out a gene ontology (GO) classification, based on the term “Endocytic Transport” revealing a total of 171 proteins. Up and downregulation were determined by calculating the ratio between protein expression values between the cell line and MCF7 (e.g. 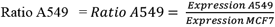). Up and down-regulated proteins are represented in green and red, respectively. A fold change ≥ 2 and ≤ 0.5 was used as the cut-off for upregulated and downregulated proteins, respectively.

Analysis of Table S1 revealed key differences in protein expression that may drive differential NP uptake rates across cell lines. The synaptosome-associated protein-23 (SNAP23), crucial for cell fusion and early endosomal function, was upregulated in A549, HeLa, and KM12C cells^17,27^. Vimentin (VIM), essential for late endocytic trafficking, showed significant upregulation in A549 and HeLa cells^28^. Unconventional myosin-Ie (MYO1E), implicated in clathrin-mediated endocytosis, and DH domain-containing protein 2 (EHD2), essential for endosomal vesicle formation, were both overexpressed in A549 and HeLa cells, suggesting a more active endocytic system^29,30^. DH domain-containing protein 2 (EHD2), another key regulator of endocytosis responsible for formation and maintenance of endosomal vesicles, was also found overexpressed in A549 and HeLa cells^30^. Caveolin-1 (CAV1) upregulation followed a pattern similar to NP uptake, highlighting its potential role in this process. As a scaffolding protein in caveolar membranes, CAV1 is essential in endocytosis and transcytosis, reinforcing the idea that caveolae-dependent mechanisms may be a key driver of Au@mSi-PEI NP internalization^31^. These protein expression trends seem to contribute to enhanced endocytic phenomenon, reflected in the faster uptake rates observed for lung and cervical cancer.

### Analysis of ECM Proteins Expression

**Table S2.** ECM-related proteins, filtered by carrying out a gene ontology (GO) classification, based on the term “Extracellular Matrix” revealing a total of 120 proteins. Up and downregulation were determined by calculating the ratio between protein expression values between the cell line and MCF7 (e.g. 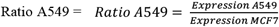). Up and down-regulated proteins are represented in green and red, respectively. A fold change ≥ 2 and ≤ 0.5 was used as the cut-off for upregulated and downregulated proteins, respectively.

| Protein | Gene | Ratio A540 | ratio HeLa | Ratio KM12C |
| --- | --- | --- | --- | --- |
| P04217 | A1BG | 1.32897 | 1.844236 |  |
| P01023 | A2M | 1.326754 | 2.529975 | 1.02563 |
| O00468 | AGRN | 1.492754 | 2.092516 | 0.866998 |
| P02765 | AHSG | 1.812283 | 2.681835 | 0.992381 |
| P04083 | ANXA1 | 220.0907 | 46.89729 | 9.390691 |
| P50995 | ANXA11 | 1.789986 | 1.137725 | 1.91124 |
| P07355 | ANXA2 | 8.844179 | 3.868842 | 2.471011 |
| P09525 | ANXA4 | 18.25557 | 0.427643 | 0.517038 |
| P08758 | ANXA5 | 1.74704 | 1.490556 | 0.567626 |
| P08133 | ANXA6 | 0.31137 | 0.68143 | 0.048384 |
| P20073 | ANXA7 | 1.193577 | 0.74 | 0.803493 |
| P02647 | APOA1 | 0.590616 | 0.745065 | 0.547634 |
| P02656 | APOC3 |  | 1.525676 | 2.213195 |
| P02649 | APOE | 0.810051 | 2.690199 | 1.34587 |
| P02749 | APOH | 2.448151 | 1.30354 | 1.294913 |
| P25311 | AZGP1 | 1.242177 | 1.951885 | 1.033316 |
| P50895 | BCAM | 0.402959 | 0.371831 | 0.057042 |
| P18075 | BMP7 |  |  |  |
| P27797 | CALR | 1.13714 | 1.785369 | 1.051865 |
| O14936 | CASK | 1.467482 | 0.743876 | 0.943096 |
| P48509 | CD151 | 3.023062 | 1.527805 | 0.841035 |
| P19022 | CDH2 | 0.931594 | 3.48766 | 1.27265 |
| Q05315 | CLC | 6.999098 | 9.952626 |  |
| P10909 | CLU | 0.987364 | 2.899027 | 0.075746 |
| Q99715 | COL12A1 | 2.112505 | 2.614408 | 1.075873 |
| P39060 | COL18A1 | 0.737116 |  |  |
| **P02452** | **COL1A1** | **2.493416** | **2.178591** | **0.910413** |
| P08123 | COL1A2 | 0.85465 | 1.133455 | 1.085136 |
| P12109 | COL6A1 | 0.991795 | 2.344587 | 4.502421 |
| P04080 | CSTB | 1.306614 | 1.106417 | 1.964935 |
| P07858 | CTSB | 2.637184 | 0.683623 | 2.242007 |
| P53634 | CTSC | 8.952306 | 14.47646 | 6.862417 |
| P07339 | CTSD | 2.091955 | 0.540813 | 0.96125 |
| P09668 | CTSH | 0.585597 | 0.691487 | 0.458323 |
| P07711 | CTSL | 3.331854 | 1.236421 | 0.353028 |
| Q9UBR2 | CTSZ | 12.16867 | 8.875278 | 1.062447 |
| Q14118 | DAG1 | 1.023319 | 1.624542 | 1.867091 |
| Q12959 | DLG1 | 0.919875 | 0.710832 | 1.044342 |
| Q9UGM3 | DMBT1 | 0.702966 | 1.294107 | 0.786838 |
| Q03001 | DST | 1.270809 | 1.686205 | 0.742201 |
| Q16610 | ECM1 | 1.718484 | 2.02337 | 0.95643 |
| P52803 | EFNA5 | 3.004466 | 1.350066 | 1.16965 |
| Q9Y5L3 | ENTPD2 | 0.442529 |  |  |
| Q96RT1 | ERBIN | 1.244011 | 0.930567 | 1.043525 |
| P23142 | FBLN1 | 3.290504 | 2.492838 | 0.558154 |
| P20930 | FLG | 0.902145 | 1.694971 | 0.776174 |
| **P02751** | **FN1** | **6.540531** | **2.065189** | **0.904584** |
| Q5SZK8 | FREM2 |  | 0.910985 |  |
| Q99988 | GDF15 |  | 1.250736 | 7.818661 |
| Q3B7J2 | GFOD2 | 0.666671 | 0.505598 | 0.236908 |
| Q92896 | GLG1 | 1.057497 | 0.594254 | 0.498737 |
| P35052 | GPC1 | 2.52571 | 2.892188 | 1.055301 |
| O75487 | GPC4 |  |  | 1.769948 |
| Q9Y625 | GPC6 | 2.068823 |  | 1.559025 |
| P51858 | HDGF | 0.880474 | 0.9663 | 0.52815 |
| P52272 | HNRNPM | 1.123168 | 0.838621 | 1.184522 |
| Q86YZ3 | HRNR | 1.73669 | 2.961116 | 0.922473 |
| Q53GQ0 | HSD17B12 | 2.464915 | 0.875046 | 4.43075 |
| P07900 | HSP90AA1 | 0.640144 | 0.56819 | 1.064083 |
| P14625 | HSP90B1 | 1.674583 | 1.060569 | 1.448033 |
| P98160 | HSPG2 | 1.423108 | 2.435391 | 0.902188 |
| Q92743 | HTRA1 | 3.244088 | 5.088924 | 1.041576 |
| P05362 | ICAM1 |  | 3.850294 | 1.063087 |
| P16144 | ITGB4 | 3.849613 | 2.386789 | 7.257904 |
| P19827 | ITIH1 |  | 0.152998 | 0.735067 |
| P19823 | ITIH2 | 1.236126 | 1.948284 | 0.735875 |
| Q14624 | ITIH4 | 1.402578 | 2.215764 | 1.029294 |
| P04264 | KRT1 | 1.313982 | 2.110487 | 1.066396 |
| P32004 | L1CAM | 0.651176 | 8.273373 | 0.336418 |
| O00515 | LAD1 |  | 3.949978 | 3.13453 |
| Q16787 | LAMA3 | 2.243738 | 1.888962 | 2.448461 |
| Q16363 | LAMA4 | 1.661494 | 1.380351 | 0.51508 |
| O15230 | LAMA5 | 2.170828 | 1.973525 | 1.659487 |
| P55268 | LAMB2 | 1.363105 | 1.150634 | 0.541437 |
| P11047 | LAMC1 | 1.254311 | 2.14197 | 3.155237 |
| **P09382** | **LGALS1** | **3.939596** | **12.28104** | **1.515778** |
| P17931 | LGALS3 | 0.720285 | 1.145415 | 0.754693 |
| Q08380 | LGALS3BP | 3.526093 | 13.34369 | 35.93923 |
| P56470 | LGALS4 | 1.899847 | 3.088118 | 4.585358 |
| P49257 | LMAN1 | 1.812742 | 2.151038 | 0.928698 |
| Q14766 | LTBP1 |  |  |  |
| P55081 | MFAP1 | 0.680849 | 1.332129 | 1.436903 |
| P14780 | MMP9 | 0.483103 | 0.773028 | 0.495551 |
| P98088 | MUC5AC | 8.543386 | 1.178325 | 1.6276 |
| Q9HC84 | MUC5B | 16.26705 | 0.604469 | 0.32214 |
| Q32P28 | P3H1 | 2.001098 | 7.539928 | 1.800384 |
| P14618 | PKM | 1.66108 | 1.603611 | 2.331335 |
| P00747 | PLG | 1.698633 | 6.701704 | 2.277162 |
| Q02809 | PLOD1 | 1.227953 | 5.168799 | 2.035406 |
| O00469 | PLOD2 | 1.194194 | 1.289466 | 0.417783 |
| O60568 | PLOD3 | 0.82119 | 1.768599 | 0.900833 |
| P07602 | PSAP | 2.2869 | 1.863548 | 2.077323 |
| Q92626 | PXDN |  | 2.423137 | 1.003163 |
| P60903 | S100A10 | 4.684369 | 1.073997 | 1.838059 |
| P26447 | S100A4 | 170.1439 | 107.6346 | 3.382575 |
| P06703 | S100A6 | 23.05123 | 19.0312 | 42.9434 |
| P31151 | S100A7 | 0.816946 | 1.320453 | 0.985287 |
| P05109 | S100A8 | 0.410868 | 0.571284 | 0.970558 |
| P06702 | S100A9 | 0.381418 | 0.494333 | 0.819706 |
| P01009 | SERPINA1 | 0.747188 | 0.946115 | 0.490431 |
| P01011 | SERPINA3 | 0.694979 | 0.527524 | 0.169386 |
| P30740 | SERPINB1 | 2.568632 | 0.88134 | 1.203641 |
| Q96P63 | SERPINB12 | 1.498947 | 2.281039 | 1.051188 |
| P35237 | SERPINB6 | 0.606569 | 0.830318 | 0.351457 |
| P50452 | SERPINB8 | 1.602912 | 1.12013 | 0.810673 |
| P50453 | SERPINB9 | 6.300039 | 2.593244 | 0.596288 |
| P01008 | SERPINC1 | 1.675814 | 0.600248 | 5.529793 |
| P50454 | SERPINH1 | 0.926076 | 2.067819 | 1.098047 |
| Q9H4F8 | SMOC1 | 5.267547 | 12.95362 | 10.00454 |
| Q9UBB9 | TFIP11 | 0.947516 | 1.13645 | 1.133983 |
| P01137 | TGFB1 | 2.116185 | 1.198644 | 2.838832 |
| Q15582 | TGFBI | 5.183256 | 13.02979 | 2.18072 |
| P21980 | TGM2 | 41.63512 | 2.646266 | 1.165121 |
| P07996 | THBS1 | 0.472234 | 0.716348 | 0.203014 |
| P35442 | THBS2 | 1.509166 | 1.536634 | 10.79358 |
| P01033 | TIMP1 | 1.588545 | 2.308944 | 2.587067 |
| P35625 | TIMP3 |  | 0.240212 |  |
| O00300 | TNFRSF11B |  | 1.597872 | 0.757092 |
| Q6EMK4 | VASN | 1.031054 | 3.621838 |  |
| **P04004** | **VTN** | **2.810338** | **14.32062** | **1.835131** |

From Table S2, we identified key proteins that may influence NP diffusion in solid tumors. Collagen Type I, (COL1A), collagen type II (COL12A1), and Fibronectin (FN1) were upregulated in A549 and HeLa spheroids, compared to KM12C and MCF7, suggesting a denser ECM that hinders NP penetration^32^. Similarly, vitronectin (VTN), an adhesive glycoprotein responsible for increased cell attachment to the ECM^33^, and fibulin-1 (FBLN1), a glycoprotein that stabilizes the ECM structure^34^, followed the same pattern. Galectin-1 (LGALS1), known for upregulating metalloproteinases (MMPs) and binding to cell surface receptors and ECM^35^, was also overexpressed, along with protein glutamine gamma-glutamyltransferase 2 (TGM2), a calcium-dependent enzyme involved in ECM cross-linking^36^. The overexpression of these proteins may contribute to a denser, more tightly packed ECM structure, potentially hindering the movement of NPs through the tumor tissue and resulting in the observed reduced penetration depths in these spheroids^2^.

### Nanoparticle Behavior Analysis – Segmentation in 3D and 2D Models

**Figure S10.**
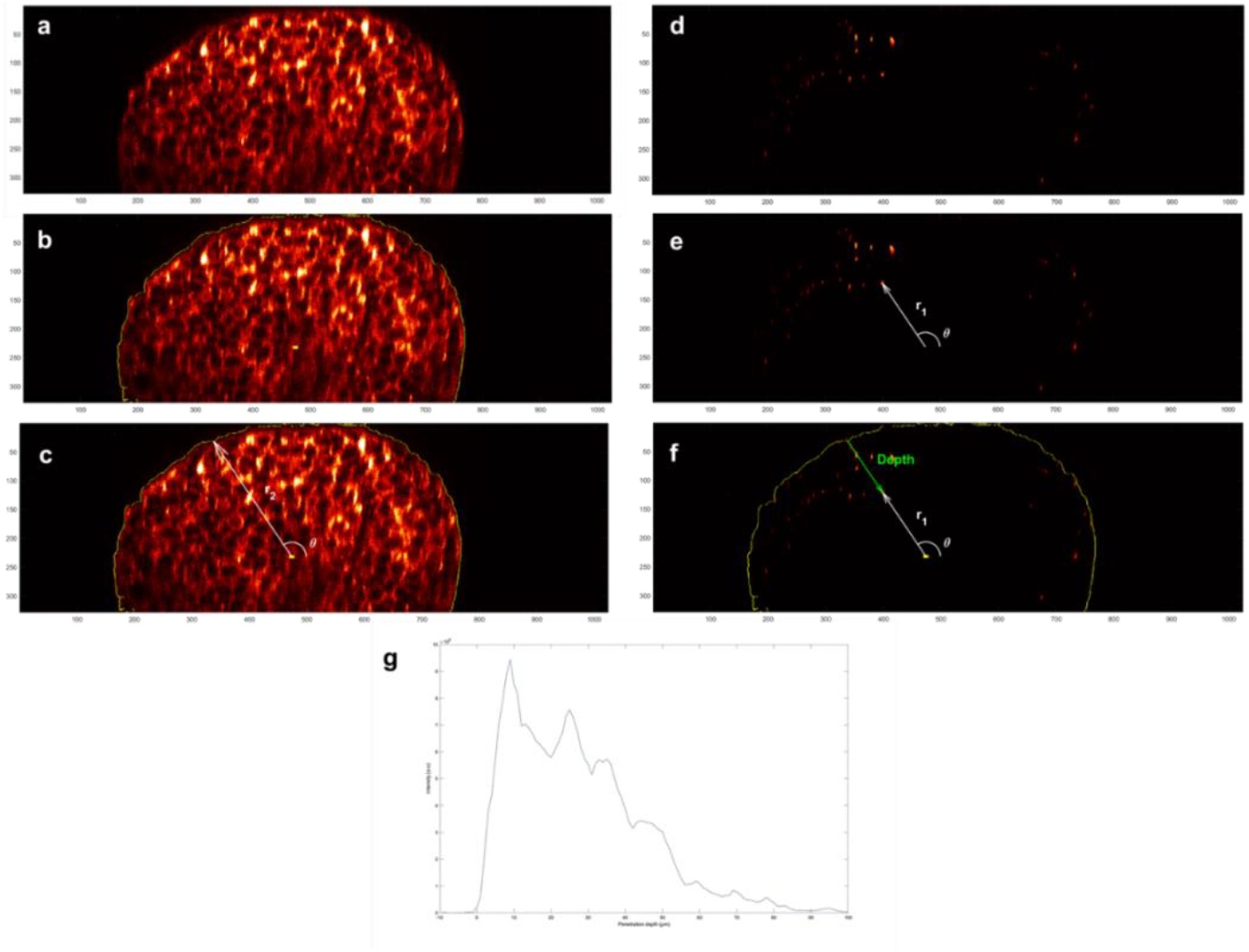
Overview of the software for NP behavior analysis in 3D spheroids. (a) Spheroid detection by the software for each xy plane of the z stack, and resulting orthogonal view of the 3D structure. (b) Preprocessing steps to determine the spheroid edge, including median filtering, noise removal, edge detection via Sobel algorithm. The x,y coordinates of the spheroid centre are determined by averaging the centroid positions of the segmented areas in 20 adjacent stacks around the plane where the spheroid appears widest. The z coordinate of the centre is taken as the z coordinate of this plane. (c) Transformation of data into ellipsoidal coordinates, storing azimuth, elevation, and distance for each point. The azimuth angle is not shown in this 2D sideview. (d) NP detection on channel 2. (e) Assignment of ellipsoidal coordinates to every pixel in the nanoparticle channel, and (f) calculation of the distance of each pixel from the spheroid edge to determine penetration depth. (e) Penetration profile from the integration of nanoparticle intensity over distance.

**Figure S11.**
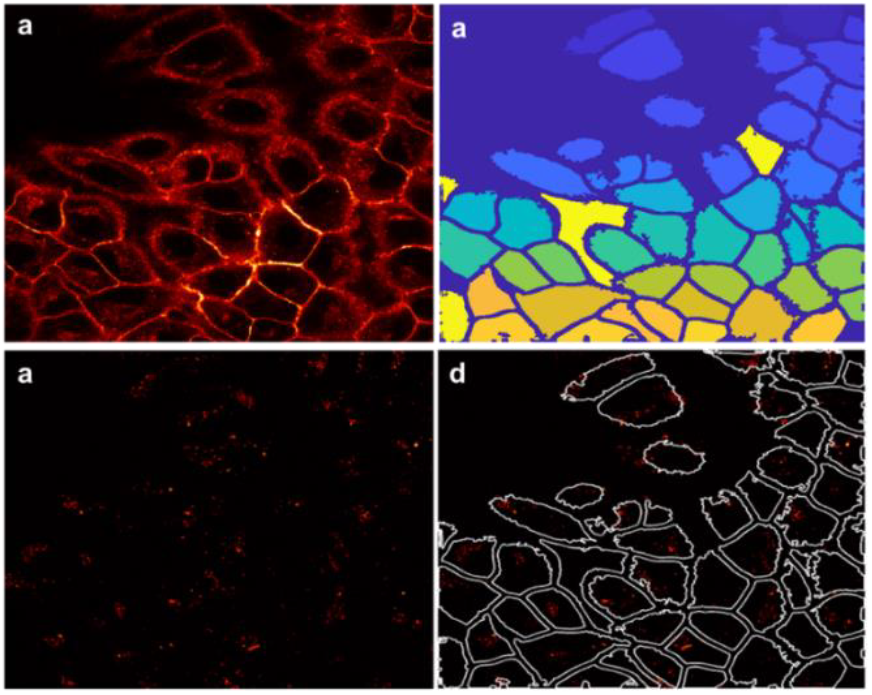
Overview of the software for NP behavior analysis in 2D cell monolayers. (a) Preprocessing of cell membrane data with median filtering. (b) Segmentation of individual cells using the Cellpose deep learning algorithm, performed on the most representative plane of the z-stack. Parameters such as cell threshold, cell diameter, and flow error tolerance are optimized for accurate segmentation. Postprocessing steps including erosion to refine segmentation boundaries. (c) NP identification in channel 2 (d) Nanoparticle intensity integration: labelled masks were used to compute nanoparticle intensity in each cell.

## Notes

### Competing Interest Statement

The authors have declared no competing interest.

